# Identifying and targeting key driver genes for collagen production within the 11q13/14 breast cancer amplicon

**DOI:** 10.1101/2024.03.27.587019

**Authors:** Daniela Araiza-Olivera, Tatiana Y. Prudnikova, Cristina Uribe-Alvarez, Kathy Q. Cai, Janusz Franco-Barraza, Jesús M. Dones, Ronald T. Raines, Jonathan Chernoff

**Affiliations:** Cancer Signaling and Microenvironment Program, Institute for Cancer Research, Fox Chase Cancer Center, Philadelphia, Pennsylvania; Histopathology Facility, Institute for Cancer Research, Fox Chase Cancer Center, Philadelphia, Pennsylvania; Marvin and Concetta Greenberg Pancreatic Cancer Institute, Institute for Cancer Research, Fox Chase Cancer Center, Philadelphia, Pennsylvania; Department of Chemistry, Massachusetts Institute of Technology, Cambridge, MA

## Abstract

Genetic studies indicate that breast cancer can be divided into several basic molecular groups. One of these groups, termed IntClust-2, is characterized by amplification of a small portion of chromosome 11 and has a median survival of only five years. Several cancer-relevant genes occupy this portion of chromosome 11, and it is thought that overexpression of a combination of driver genes in this region is responsible for the poor outcome of women in this group. In this study we used a gene editing method to knock out, one by one, each of 198 genes that are located within the amplified region of chromosome 11 and determined how much each of these genes contributed to the survival of breast cancer cells. In addition to well-known drivers such as *CCND1* and *PAK1*, we identified two different genes (*SERPINH1* and *P4HA3*), that encode proteins involved in collagen synthesis and organization. Using both *in vitro* and *in vivo* functional analyses, we determined that *P4HA3* and/or *SERPINH1* provide a critical driver function on IntClust-2 basic processes, such as viability, proliferation, and migration. Inhibiting these enzymes via genetic or pharmacologic means reduced collagen synthesis and impeded oncogenic signaling transduction in cell culture models, and a small-molecule inhibitor of P4HA3 was effective in treating 11q13 tumor growth in an animal model. As collagen has a well-known association with tissue stiffness and aggressive forms of breast cancer, we believe that the two genes we identified provide an opportunity for a new therapeutic strategy in IntClust-2 breast cancers.

## Introduction

Breast cancer encompasses diverse entities with distinct clinical, morphological, and molecular features. Current classification relies on morphology (histological type and grade) and key markers like estrogen receptor (ER) and human epidermal growth factor receptor 2 (HER2) (1). This classification influences prognosis and guides therapeutic strategies (2, 3).

In recent years, advanced molecular analyses have greatly enhanced our understanding of breast cancer mutations and genetic alterations, leading to a more precise classification.

Breast cancers frequently display regions of chromosomal amplification and deletion. While such chromosomal aberrations are common in breast cancer, identifying their precise nature, relevance to tumor initiation and progression, and potential for therapeutic targeting remains a challenge. In addition to point mutations and small-scale structural changes, breast cancer genomes frequently exhibit significant structural alterations, including large-scale gains and losses of genetic material, known as copy number alterations (CNAs) (3, 4). Recurrent patterns of CNAs suggest their causal involvement in tumorigenesis. Recent studies have focused on somatic alterations in breast cancer, including mutations and CNAs (1, 4, 5). Furthermore, transcriptomic, and genomic data have been integrated to link recurrence with deregulated gene expression (4-6). The Molecular Taxonomy of Breast Cancer International Consortium analyzed the genomic and expression profiles of 2,000 breast tumors (3, 5) revealing that recurrent and focal copy number alterations (loss or gain) affect the expression of genes within these regions, including amplifications mapping to *ERRB2, MYC, CCND1, CCNE1, MDM2* and *MDM4* and deletions of *PTEN, PPP2R2A* and *MAP2K4* (7).

Using comprehensive genomic, transcriptomic, and clinical annotations from this large sample collection, breast tumors have been classified into 10 subtypes based on patterns of CNAs and gene expression (1, 3, 8). This classification system, known as Integration Clusters (IntClust), offers a genome-wide driver-based classification for breast cancer patients, correlating tumor genomes and transcriptome data for clinical purposes (2-4, 9, 10).

One such cluster, termed IntClust-2, is defined by a recurrent amplification of regions of 11q13/14, which occur in up to 10% of breast cancers and are associated with complete resistance to chemotherapy and represent the worst prognosis of all ER-positive tumors (10-year disease-specific survival rate ∼50%) (1, 3, 5, 9, 11).

A key challenge in analyzing CNAs is determining the relative cancer contributions of amplified genes within these regions. For instance, in the *HER2* amplicon on 17q12 (IntClust-5 tumors), most or all the oncogenic activity is thought to be associated with a single gene (*viz, HER2* itself); whereas in other cases ensembles of genes likely make independent contributions to oncogenesis (3, 4). This scenario is particularly germane to the 11q13/14 amplicon(s), which, in breast cancer, can be subdivided into several distinct regions. In some cases, the entire region is amplified; in others, only one or a few of these regions is amplified. These data indicate that chromosome region 11q13 is rich in potential oncogenes that are relevant to breast cancer (1, 7, 9).

*CCND1*, encoding cyclin D, has received the most attention as “the” oncogene in the 11q13/14 region. Amplification of *CCND1* and elevated cyclin D expression are common in breast cancer, correlating strongly with increased cell proliferation (12). However, *CCND1* represents at most only a fraction of the oncogenic stimulus in cells bearing chromosome 11 CNAs, as many breast cancer cells show a “firestorm” pattern of amplification of other nearby gene regions, often excluding *CCND1* (13). Indeed, in the seminal study of Curtis *et al*., which defined Integration Clusters in breast cancer, showed that the IntClust-2 group, containing amplified *PAK1* and adjacent genes at 11q13.5, was associated with aggressive disease and poor survival (3, 5). These tumors, predominantly luminal A or B type and varying in the estrogen-receptor (ER) status (7), were associated with one of the worst prognoses among the ten IntClust categories (4, 11). Interestingly, the Hahn group, in an unbiased search for protein kinases that transform breast epithelial cells, reported that *PAK1* exerted a powerful effect on the acquisition of anchorage-independence and other hallmark properties of transformed cells (14). Thus, the amplicon in question may contain at least two important cancer driver genes (*i*.*e*., *CCND1* and *PAK1*), and quite possibly more.

While *CCND1* and *PAK1* gene amplification likely contribute to the pathogenesis of IntClust-2 breast cancers, the roles of other potential oncogenes in the neighboring amplified region of chromosome 11 remain unclear. What has been missing to date is a comprehensive screen to identify and quantitate the relative contributions of each of these amplified genes. An approach is important to identify gene products that might provide suitable targets for therapeutic intervention, either alone or in combination with targeted inhibitors of established 11q13/14-encoded oncoproteins such as CyclinD/Cdk4/6 (*e*.*g*. palbociclib) and/or Pak1 (*e*.*g*. G5555) (15). Recently, RNAi methods have been applied to identify such additional drivers. For example, Choi *et al*. used RNAi against six candidate oncogenes in the 11q13.5 cluster to assess their contribution to paclitaxel sensitivity in OVCAR-3 ovarian cancer cells (16). This study yielded *RSF1* as a strong candidate, but similar studies have come to different conclusions (17). The lack of consistency may result from the use of different cell lines and technical issues inherent in RNAi.

Recently, gene-editing technologies have been developed and are potentially suitable for identifying oncogenic drivers. The most facile of these is the CRISPR-Cas9 system, which has been adapted to induce double-stranded breaks in genomic DNA in mammalian cells (18, 19). Repair of these CRISPR-Cas9 induced lesions by homologous recombination pathways often results in small insertions or deletions that disrupt the reading frame of the encoded protein (19). Because of its ease of use, large-scale high-throughput CRISPR-Cas9 screens have been instrumental in identifying oncogenic drivers and tumor suppressors across various systems (18, 20). Here, we use a CRISPR-Cas9-based gene knock out approach to probe the oncogenic contribution of each of 198 genes between *CCND1* and *PAK1* on chromosome 11 in breast cancer cell lines. We found two previously understudied genes – the collagen prolyl hydroxylase *P4HA3* and the collagen chaperone *SERPINH1* – among the top contributing genes to the proliferation of several IntClust-2 cell lines. As will be discussed, these findings offer a potential new therapeutic strategy in this category of breast cancer.

## Materials and Methods

### Reagents

P4HA3 inhibitors: ethyl 3,4-dihydroxybenzoate (#HY-W016409) from MedChemExpress and diethyl pythiDC synthesized as described previously (21) or purchased from MedKoo Biosciences, Inc (#563602).; SERPINH1: Col003 (#HY-124817) from MedChemExpress.

Transfection reagents: P4HA3 (#L-008479-01-0005), SERPINH1 (#L-011230-00-0005) and DharmaFECT (#T-2001-01) were purchased from Dharmacon Reagents (Horizon).

### Cell Culture

Human breast cancer cell lines HS578T, MDA-MB-436, MDA-MB-361 and ZR-75-1 were obtained from the ATCC (Manassas, Virginia). MDA-MB-231 and HCC1395 were generously provided by Neil Johnson (Fox Chase Cancer Center, PA). HS578T and MDA-MB-436 were adapted and maintained in Dulbecco’s Modified Eagle’s Medium (DMEM) (Sigma) supplemented with 10% fetal bovine serum (FBS), 1.5 g/L NaHCO_3_, 5% L-glutamine,1% antibiotics (penicillin/streptomycin) and 0.1 unit/ml bovine insulin. HCC1395, MDA-MB-231 and ZR-75-1 were adapted and maintained in RPMI 1640 culture medium (Sigma) supplemented with 10% FBS, 1.5 g/L NaHCO_3_, 5% L-glutamine,1% antibiotics (penicillin/streptomycin). HCC1395 also required 4.5 g/L glucose, 10 mM HEPES and 1 mM pyruvate. MDA-MB-361 was maintained in 1:1 DMEM/Leibovitz’s L-15 (Sigma) supplemented with 10% fetal bovine serum (FBS), 1.5 g/L NaHCO_3_, 5% L-glutamine,1% antibiotics (penicillin/streptomycin). Non tumor MCF-10 cells were grown in DMEM with 10% fetal bovine serum (FBS), 1.5 g/L NaHCO_3_, 5% L-glutamine,1% antibiotics (penicillin/streptomycin). All cell lines were cultured in a humidified incubator with 5% CO_2_ at 37° C and evaluated for mycoplasma.

### CRISPR screen

HEK-293T cell line was transduced with a Cas9 lentiviral vector with blasticidin selection marker (Addgene# 52962) and selected with blasticidin (8 ug/ml). Single clones with high Cas9-expression were established and used in the CRISPR screen as described by Wang et al. (22).

A CRISPR library was constructed to target 198 coding genes located on 11q13/14 between *CCND1* and *GAB2* genes (6 sgRNAs per gene), as well as non-targeting sgRNAs and known essential control genes. In brief, oligos were synthesized (IDT), and ligated into pLenti-sgRNA vector (Addgene 71409) with puromycin selection marker, containing an U6 promoter to drive sgRNA expression. The plasmid libraries were then transfected into Lenti-X cells (Takara) to produce lentiviral pools, which were subsequently transduced into SUM52 breast cancer cell line. Cells were infected with the libraries at a multiplicity of infection of 0.3–0.5, and after 48 h were selected with puromycin (1 μg ml−*1*) for 3–5 days until control cells without virus were dead. Cells were expanded for another 2–3 days and aliquots were saved as D0 stocks. The remaining cells cultured for another 14 days and collected for further processing. To maintain library complexity, the screens were performed at ∼500 × cell number coverage per sgRNA (∼3 million cells were transduced). All screens were performed in biological replicate.

DNA was extracted from each sample using DNA Mini Kit (Qiagen) with the manufacturer’s protocol. 3 μg of genomic DNA per sample was used as template for PCR to amplify the sgRNA region using Ex Taq DNA polymerase (Takara). Next Generation Sequencing (NGS) was performed on an Illumina HiSeq (Illumina) to determine the abundance of sgRNA.

To analyze the samples, the average log2 fold change of all target sgRNAs for each gene (gene score) was calculated. To compare between samples, the difference in gene scores for D0 and D14 was calculated to identify differentially scoring genes.

1. Primer Sequences for sgRNA Quantification Forward: AATGATACGGCGACCACCGAGATCTACACGAATACTGCCATTTGTCTCAAGATCTA Reverse: CAAGCAGAAGACGGCATACGAGATCnnnnnnTTTCTTGGGTAGTTTGCAGTTTT (nnnnnn denotes the sample barcode)
2. Illumina Sequencing Primer CGGTGCCACTTTTTCAAGTTGATAACGGACTAGCCTTATTTTAACTTGCTATTTCTAGCTCTAAAAC.
3. Illumina Indexing Primer TTTCAAGTTACGGTAAGCATATGATAGTCCATTTTAAAACATAATTTTAAAACTGCAAACTACCCAAGAAA.

### siRNA transfection

All the breast cell lines were seeded in a 96-well plate at 5000 cells/per well and cultured in the corresponding antibiotic-free medium 24 h before the experiment. For the transfection it was used 25 nM of P4HA3 or SERPINH1 siRNA using the cationic lipid Dharma*FECT* 1 (0.20 μL/well). The cells were incubated at 37°C in 5% CO_2_ for 48 h for the proper analysis. (According to the manufacturer’s protocol).

### Cell viability

Breast tumoral and non-tumoral cell lines were seeded in 96-well plates at 5000 cells/well in the corresponding medium and incubated overnight. After 24 h, the supernatant was removed and fresh medium with siRNA or increasing concentrations of P4HA3 and SERPINH1 inhibitors (0, 0.2, 0.4, 0.8, 2, 4, 8 and 20 μM) were respectively added. Cell viability was evaluated after incubation with drugs using Alamar Blue fluorescent assay (Life Technologies). Experiments were done in 48 h and 0.1% DMSO was used as negative control. IC_50_ was calculated from three independent experiments.

### Proliferation assays

For this experiment 100,000 cells in a 6-well plate were seeded and incubated overnight. After 24 h the respective P4HA3 and SERPINH1 siRNA (25 nM) or inhibitor (2 μM) was added. Cells were trypsinized with Tryp-LE express (Gibco), centrifuged, resuspended in medium and counted at different incubation times (0, 24, 48, 72 and 96 h) on a TC 20 automated cell counter (Biorad). The experiments were conducted in triplicate for each timepoint. The percentage of cell proliferation was calculated by comparing it with control cells. The results are shown as mean ± standard deviation (SD) (* p < 0.05, ** p < 0.01, *** p < 0.001 *versus* control).

### Cell Migration Assay (Wound Healing)

Breast cancer cells were seeded in six-well plates until 90–100% confluence was reached and manually scraped with a 200 μL pipette tip. The cells were washed and then grown in their respective media containing siRNA (25 nM) or small molecules inhibitors (2 μM).

Scratched areas were monitored at 0, 24 and 48 h using an EVOS fluorescence microscope. Images were acquired at 100× magnification and analyzed by the number of cells that cross into the wound area from their reference point at time zero. Each image was made in triplicate and quantified using ImageJ software (National Institute of Mental Health, Bethesda, Maryland, USA) to determine the migration rate of the cells. The percentage of migration was quantified by measuring the size of the cell free-area using the formula: %Cell Migration = ((area0HR - area24HR)/ area0HR) *100. Prism 7.0 (GraphPad, San Diego, CA) was used for data analysis that represent the mean (SD) of three independent experiments. (* p < 0.05, ** p < 0.01 *versus* Control).

### Western Blot

Each cell line was grown in six-well plates until 80% confluence was reached and incubated for 48 h without or with the respective treatment (siRNA or inhibitor). For the protein extraction, RIPA buffer, protease, and phosphatase cocktail inhibitors (Sigma Aldrich) were used to scrape the cells. The cells were centrifuged for 15 min at 15,000 x g at 4°C and the supernatant was quantified with Quick Start Bradford 1X Dye Reagent (BioRad), using an albumin curve to determine the concentration of protein to be placed. Laemmli Buffer (BioRad) and 10% β-mercaptoethanol were added to the samples and boiled for 5 minutes. The protein was loaded in a 12% SDS–PAGE gel and transferred to a PVDF membrane. It was blocked for 45 min with a 5% milk-0.1% TBS-Tween 20 (blocking solution). A dilution of 1:1000 of the primary antibody in the blocking solution 4°C was done overnight. The membrane was washed 3 times with 0.1% TBS-Tween 20 every 10 min and incubated with the secondary antibody at a 1:10000 dilution in 0.1% TBS-Tween 20 for 1 h at room temperature. For the development, isovolumetric amounts of ECL reagent were added. The chemiluminescence was detected on an X-ray, which was developed and fixed.

Primary antibodies used in this study were: Collagen I (Proteintech #14695-1-AP), Collagen IV (Invitrogen #PA5-104508), pDDR1 (Tyr513) (Cell Signaling Tech. #14531), DDR1 (D1G6) (Cell Signaling Tech. #5583), P4HA3 (Proteintech #23185-1-AP), SERPINH1 (Biorad #VMA00970), phospho-ERK1/2 (Cell Signaling Tech pThr202/pTyr204) (#9101), -total ERK1/2 (Cell Signaling Tech #9102), -GAPDH (#4138).

Secondary antibodies were: Peroxidase AffiniPure Goat Anti-Mouse IgG (H+L) (# 115-035-003) and Peroxidase AffiniPure Goat Anti-Rabbit IgG (H+L) (# 111-035-003) from Jackson ImmunoResearch Labs.

### Xenografts

Female 6-week-old *nu/nu* mice were injected subcutaneously into the flank with 10^6^ breast cancer cell lines (HS578T or ZR-751) in 0.1 ml 30% Matrigel (BD Biosciences)/PBS. When tumor volume reached 100 mm^3^ measured with vernier calipers and tumor volume was calculated by (length x width2) x 0.5), mice were randomly divided into control and treated groups per cell line (8 mice in each group). Diethyl pythiDC was dissolved in 5% DMSO, 40% PEG 300 and 5% Tween 80 and 50% saline solution then administrated via intraperitoneal injection at 100 mg/kg per week. Tumor growth and weight of the animals were monitored every week. Mice were euthanized after 4 weeks of treatment. Tumors were resected, individual portions of tumors were snap-frozen in liquid nitrogen for preparation of protein lysates and fixed in 10% formalin and paraffin embedded for immunohistochemical studies. A two-sided Wilcoxon rank sum test (α = 0.05) to compare tumor volume at 4 weeks between treated and control mice will be used. With 8 mice per group, we will have 80% power to detect a between-group difference in mean tumor volumes.

All animal procedures were performed according to the approved protocols of the Fox Chase Cancer Center Institutional Animal Care and Use Committee.

### Immunohistochemistry

Collected tumors fixed in 10% formalin and embedded in paraffin were used for immunohistochemical studies. Slides were stained with Ki-67, Caspase-3 and Trichrome for collagen and later scanned using an Aperio ScanScope CS 5 slide scanner (Aperio, Vista, CA, USA). Scanned images were then viewed with Aperio’s image viewer software (ImageScope, version 11.1.2.760, Aperio). A pathologist manually outlined selected regions of interest. The immune cells quantifications were performed with Aperio V9 or positive pixel count (PPC) algorithm.

### Colorimetric Assay

Breast cancer cells (1×10^5^) were seeded into 24-well plates and incubated for 48 h in the presence or absence of the small molecule inhibitors. Quantification of total secreted collagen in the cell culture supernatant was performed using sirius red collagen detection kit (9062, Chondrex). The Collagen 1A levels were measured using the COL1A1 Elisa kit (MBS703198, MYBioSource.com). Internal and external amount of total collagen was also detected via hydroxyproline using Hydroxyproline assay kit (6017, Chondrex). All colorimetric assays were performed according to the manufacturer’s protocol.

### Statistical Analysis

The data were analyzed and plotted using GraphPad Prism software version 7.00 for Windows (GraphPad Software, California USA). The statistical analysis was performed through the one-way or two-way ANOVA test of the Tukey, Dunnett, or Sidak multiple comparisons tests. All values reflect the mean (SD), with a significance cut-off of * p < 0.05, ** p < 0.01, *** p < 0.001.

### Data availability

The data analyzed in this study were obtained from The Cancer Genome Atlas (TCGA) database downloaded from cBioportal and the Broad Institute’s Cancer Dependency Map project.

The data generated in this study are available upon request from the corresponding author.

## Results

Breast cancer is caused by a set of mutations and other genetic alterations including the amplification of a small region of the chromosome 11q13 (IntClust-2) (**Fig. 1A**). To determine the potential oncogenes that are relevant for breast cancer, a CRISPR-Cas9 systematical deletion of all genes within this region was performed. For our comprehensive screen to identify all driver genes in the IntClust-2 amplicon, we constructed six separate sgRNA lentiviral plasmids against each of the 198 genes in this region, spanning from *CCND1* at 11q13.3 to *GAB2* at 11q14.1, and used these in a pooled CRISPR-Cas “drop-out” screen to assess the relative contributions of each gene to the proliferation of an IntClust-2 (SUM52, >5 copies of *PAK1* in the 11q13 region) *vs*. a non-IntClust-2 (ZR75-1) breast cancer cell line (**Fig. 1B**). We found ten strong candidates (*i*.*e*. depleted sgRNAs, indicating target genes that consistently contributed to the survival and/or proliferation of the IntClust-2 breast cancer cell line SUM52) (**Fig. 1C** and **Supplemental Table 1**). As expected, *CCND1* and *PAK1* appeared, but we also found a few genes that had not been previously considered, that are specifically required for collagen synthesis and processing (23). We focused on two strong “hits” in this screen, *P4HA3* and *SERPINH1* (*a*.*k*.*a. HSP47*), as they encode proteins with a related function, namely, collagen synthesis.

**Figure 1.**
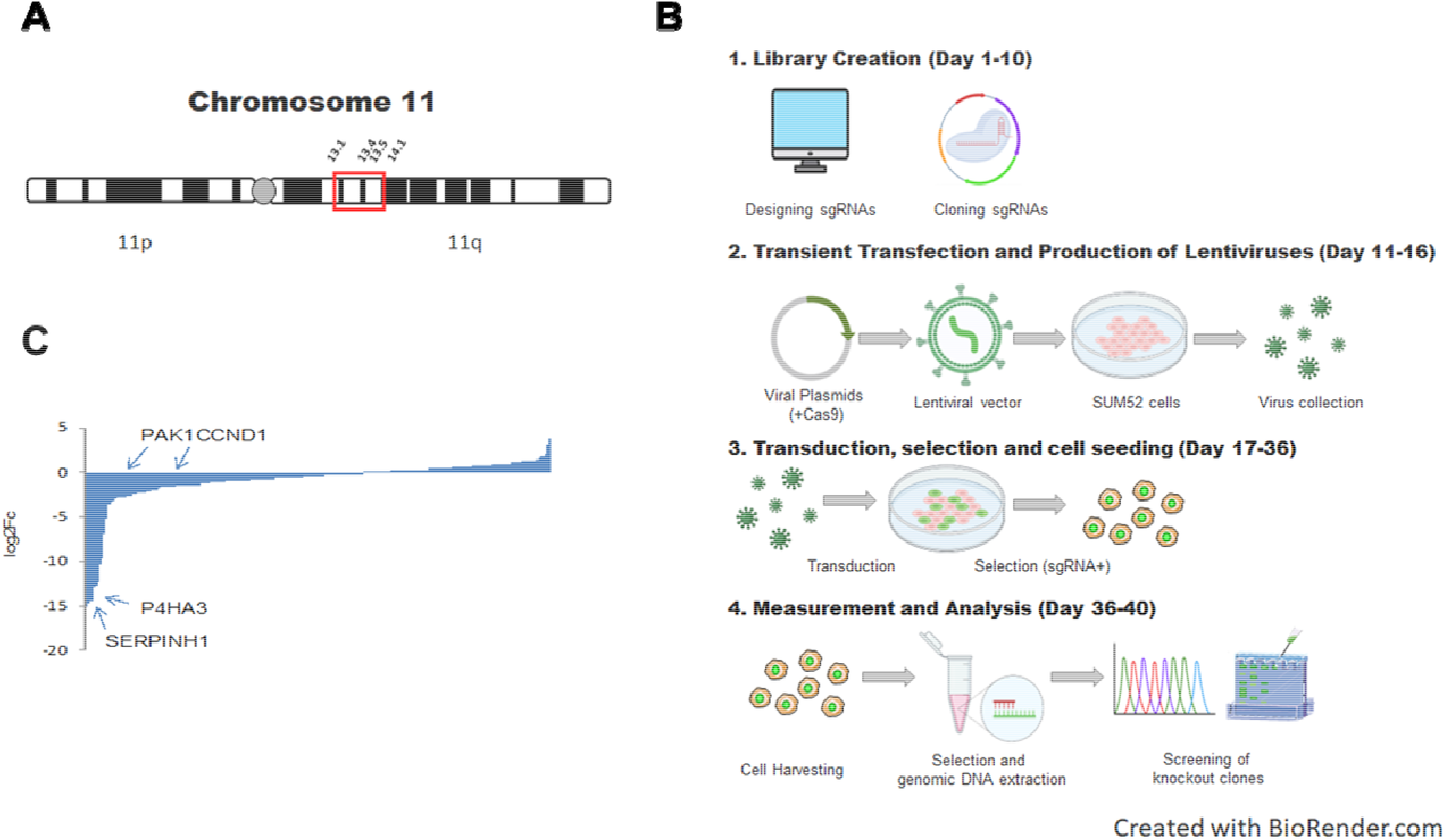
CRISPR/Cas screen for driver genes in 11q13. **A**) Structure of 11q13/14 amplicon in breast cancer highlighted in red rectangle. **B**) Primary screen: Relative roles of 198 genes within 11q13 to survival of SUM52 cells. **C**) Prominent “hits” in the IntClust-2 amplicon.

### IntClust-2 cells overproduce collagen and activate collagen signaling pathways

To analyze if our findings apply to breast cancer in general or just to IntClust-2 tumors, we chose a panel of seven different cell lines representing IntClust-2 (HS578T, MDA-MB-436 and MDA-MB-231), non-IntClust-2 (HCC1395, MDA-MB-361 and ZR-751) and a control (MCF-10). Table 1 shows that all the IntClust-2 cell lines have higher gene expression of SERPINH1 and P4HA3 compared to the non-IntClust-2. (**Table 1**).

**Table 1.** Genetic profiles of breast cancer cell lines. Gene expression and classification of the cell lines based on the amplicon obtained from DepMap.

| **Cell line** | **SERPINH1**<br>**(RNA seq)** | **P4HA3**<br>**(RNA seq)** |
| --- | --- | --- |
| HS578T | 11.3 | 2.6 |
| HCC1395 | 8.8 | 2.3 |
| MDA-MB-436 | 7.79 | 1.6 |
| MDA-MB-231 | 7.7 | 0.6 |
| MDA-MB-361 | 6.9 | 0.8 |
| ZR-751 | 5.3 | 0.05 |
| MCF-10 |  |  |

Our theory predicts an association between the IntClust-2 genomic classification and collagen-stimulated signaling in tumor cells. To address these issues, we examined if tumor cells bearing 11q13/14 amplifications express increased amounts of SERPINH1, P4HA3, collagen and enhanced DDR signaling. Figure 2 shows that IntClust-2 protein levels were increased in general compared to the non-IntClust-2.

**Figure 2.**
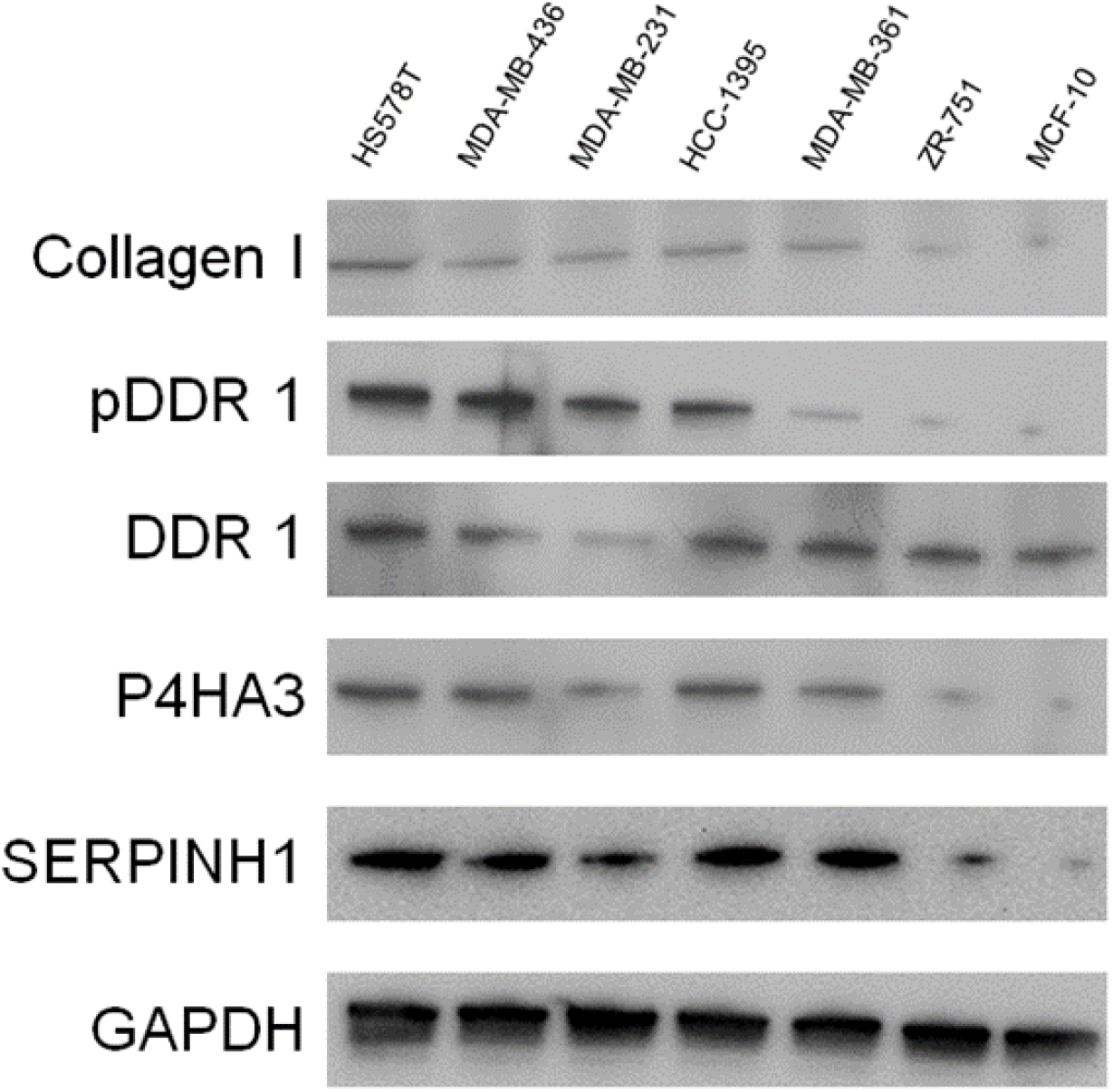
Protein levels of collagen pathway elements. Immunoblot in IntClust-2, non-IntClust-2, and control breast cancer cell lines.

### Impact of P4HA3 and SERPINH1 knockdown on cell toxicity, proliferation, migration and signaling pathways

Previously we showed that IntClust-2 cells overproduce collagen and activate collagen-stimulated signal transduction pathways (**Fig. 2**). We conducted *in vitro* functional analyses using gene knock-down to determine if the SERPINH1 and/or P4HA3 provide a critical driver action in this class of breast cancer. First, we probed that siRNA-mediated knockdown of either *P4HA3* or *SERPINH1* decreased viability around 50% of all the IntClust-2 cell lines but almost all non-IntClust-2 cell lines remained viable (**Fig. 3A**). Further analysis revealed that the siRNA led to a clear significant difference in proliferation of two IntClust-2 cell lines, but not in three non-IntClust-2 cell lines (**Fig. 3B**).

**Fig. 3.**
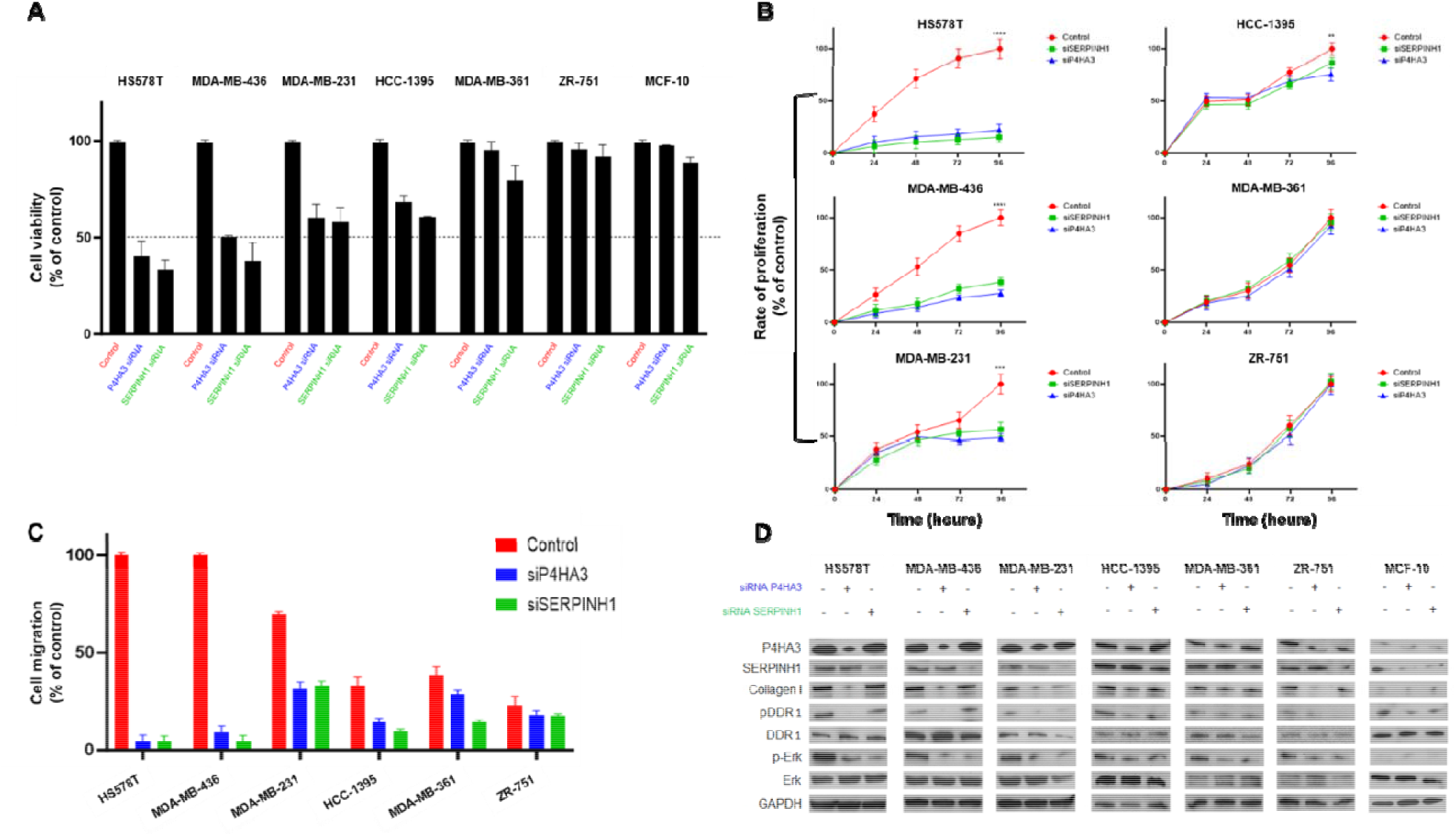
Selective role of collagen synthesis enzymes on critical functions. IntClust-2 and non-IntClust-2 cells were grown under standard conditions and treated with either 25 nM P4HA3 (blue) or SERPINH1 (green) siRNA. Control without treatment (red). Effect of P4HA3 or SERPINH1 knockdown on: **A**) Viability: Percentage of viable cells was measured after 48 h. **B**) Proliferation: Cells with the respective siRNA were counted for 0, 24, 48, 72 and 96 h. **C**) Migration: Motility was imaged at 0 and 48 h. and **D**) Signaling: Activity of P4HA3 or SERPINH1 and related pathways were quantified by immunoblot. The data percentage is the result of three independent experiments. The results are shown as mean ± standard deviation (* p < 0.05, ** p < 0.01, *** p < 0.001, \*\*\*\**p* < 0.0001 *versus* control).

Regarding the motility we observed that in general the IntClust-2 cell lines seem to migrate more than the non-IntClust-2. The knockdown of the genes prevents the healing of the scratch of the IntClust-2 and barely affects the migration of two non-IntClust-2 cells (**Fig. 3C** and **Supplemental Fig. 1**).

Experiments using siRNA show the relationship between P4HA3 or SERPINH1 respectively and collagen, DDR1 and ERK activity in the seven different cell lines (**Fig. 3D**). Through Western-blot analysis we observed that knocking down P4HA3 results in a decrease of P4HA3, collagen, pDDR1 and pERK levels in all the IntClust-2 cell lines. In the case of the SERPINH1 knockdown, just the levels of SERPINH1 and pERK were abolished. Protein levels in the non-IntClust-2 and control remained the same.

### Effect of collagen small molecule inhibitors on cellular functions

Having established the effects of P4HA3 and SERPINH1 knockdown on different cell processes, we decided to study the impact of small molecule inhibitors (**Supplemental Fig. 2**) on breast cancer cell lines. Proliferation, survival, and motility assays were assessed to detect the consequences of inhibitors related to collagen synthesis.

The cytotoxic effects of P4HA3 (ethyl 3,4-dihydroxybenzoate and diethyl pythiDC) and SERPINH1 (Col003) inhibitors were first determined by a survival assay of the cell lines when exposed to increasing concentrations (0, 0.2, 0.4, 0.8, 2, 4, 8 and 20 μM) of the drugs.

Low concentrations (less than 1.5 μM) of Col003 decreased to 50% the viability on IntClust-2 cell lines. More than 7 μM were needed to affect the viability of almost all the non-Intclust-2 cells, as well as for the control. Ethyl 3,4-dihydroxybenzoate (ethyl 3,4-DHB) reduced the survival percentage of the IntClust-2 cells with small concentrations. Non-IntClust-2 cells were also impacted with slightly higher concentrations of the SERPINH1 inhibitor. Finally, the cell viability dropped below 50% with diethyl pythiDC concentrations under 0.7 μM in the IntClust-2 cells. Up to 70% of survival was reached with the highest concentrations of diethyl pythiDC in the non-IntClust-2 and control cell lines (**Fig. 4B**). The calculated IC_50_ values for each cell line treated with the tested drugs are shown in **Supplemental Table 2**.

**Fig. 4.**
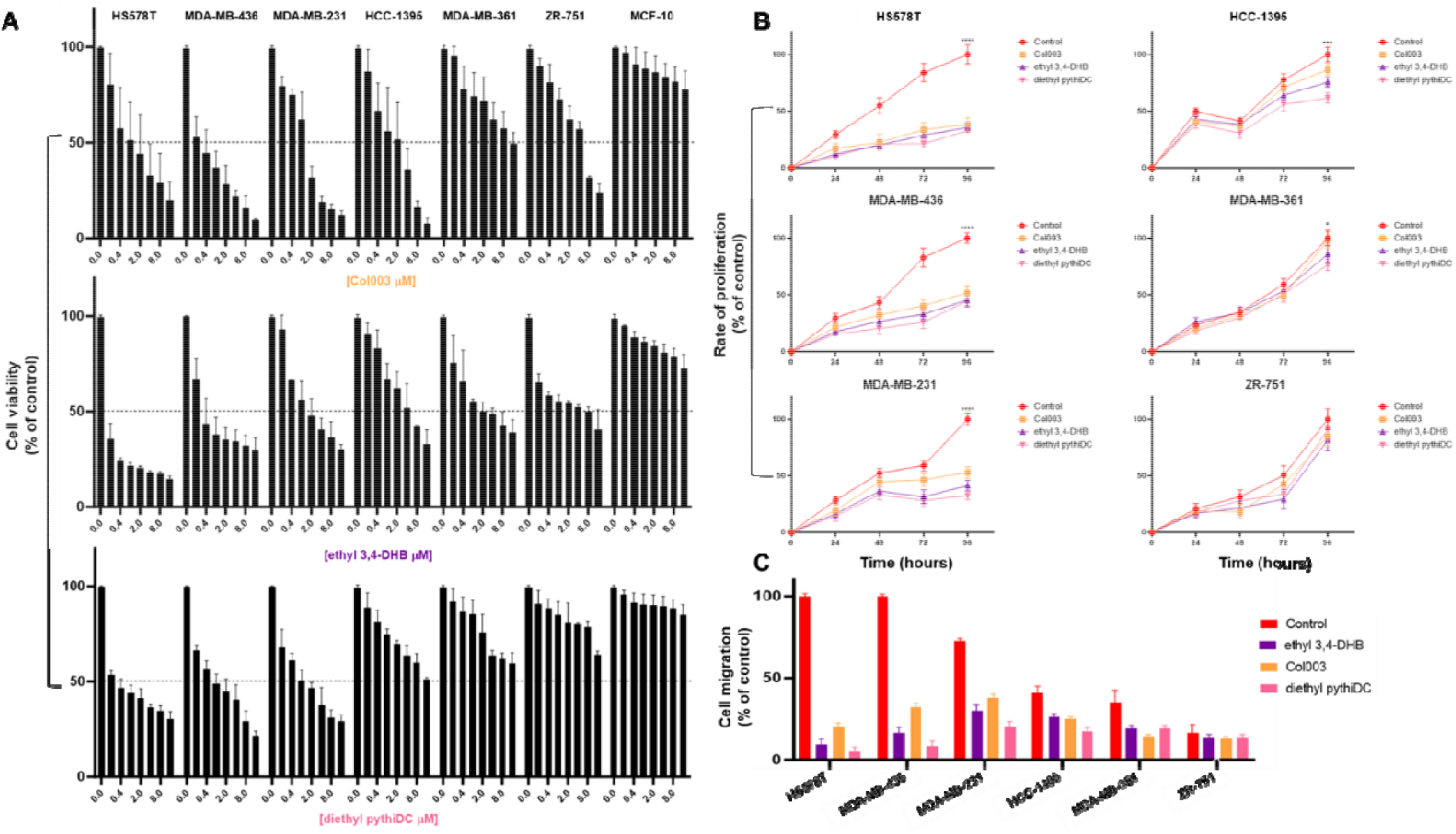
Targeted P4HA3 or SERPINH1 inhibitors effects on cell processes. All assays were performed on IntClust-2 or non-IntClust-2 cells without treatment (red) or with Col003 (orange), ethyl 3,4-DHB (purple) and diethyl pythiDC (pink). **A)** Viability: The cell lines were treated for 48 h with concentrations ranging from 0.2 to 20 μM and quantified. **B)** Proliferation: Cells treated with 2 μM of each drug were counted at different times (0, 24, 48, 72 and 96 h). **C)** Migration: A wound-healing assay was performed in the presence of 2 μM of the respective drug and images were taken during 0 and 48 h. The data percentage is the result of three independent experiments. The results are shown as mean ± standard deviation (* p < 0.05, ** p < 0.01, *** p < 0.001, \*\*\*\**p* < 0.0001 *versus* control).

Based on the genotype differences of the cell lines we studied how the P4HA3 and SERPINH1 affected proliferation. IntClust-2 and non-IntClust-2 cell lines tested had similar basal rates of proliferation. Treatment of the IntClust-2 cell lines with all the drugs decreased the cell proliferation, but the percentage difference was more evident in HS578T and MD-MB-436. This pattern of response was reversed in some of the non-IntClust-2 cell lines when the P4HA3 and SERPINH1 inhibitors were used. Only HCC1395 showed an intermediate response to the drugs (**Fig. 4B, C**), as well as with the siRNA **(Fig. 3A-C)**. Interestingly, this non-IntClust-2 cell line has elevated expression levels for P4HA3 and SERPINH1 (**Fig. 2 and Fig. 3D**).

P4HA3 and SERPINH1 are enzymes required for collagen synthesis and it is well known that collagen is associated with enhanced cell motility. We used a wound-healing assay to test if the inhibitors differentially affect the migration of the cell lines depending on their genomically-clustered group. Consistent with our previous migration results regarding the siRNA knockdown, we observed that IntClust-2 cells have a higher migration percentage compared to the non-IntClust-2. The migration of all IntClust-2 was severely decreased by the presence of all the drugs, however the P4HA3 inhibitors were more potent. Two of the non-IntClust-2 partially reduced their migration, but a lack of difference was observed in the ZR-751cells (**Fig. 4D** and **Supplemental Fig. 3**).

### IntClust-2 cells are sensitive to inhibitors of collagen synthesis

We have already shown that tumor cells bearing 11q13 amplifications present increased amounts of collagen and its dependence on P4HA3 and SERPINH1 expression. The activity of these proteins has been compromised by the presence of small molecules that inhibit collagen prolyl hydroxylases (ethyl 3,4-DHB and diethyl pythiDC) or Hsp47 (Col003). The impact of the inhibitors on the expression of total collagen, hydroxyproline and collagen I was measured in cells and in the media. Internal and external collagen I production of IntClust-2 cells decreased between 10 to 40% in the presence of diethyl pythiDC and Col003. Non-IntClust-2 cells reduction of collagen I was less than 5% (**Fig. 5A**).

**Fig. 5.**
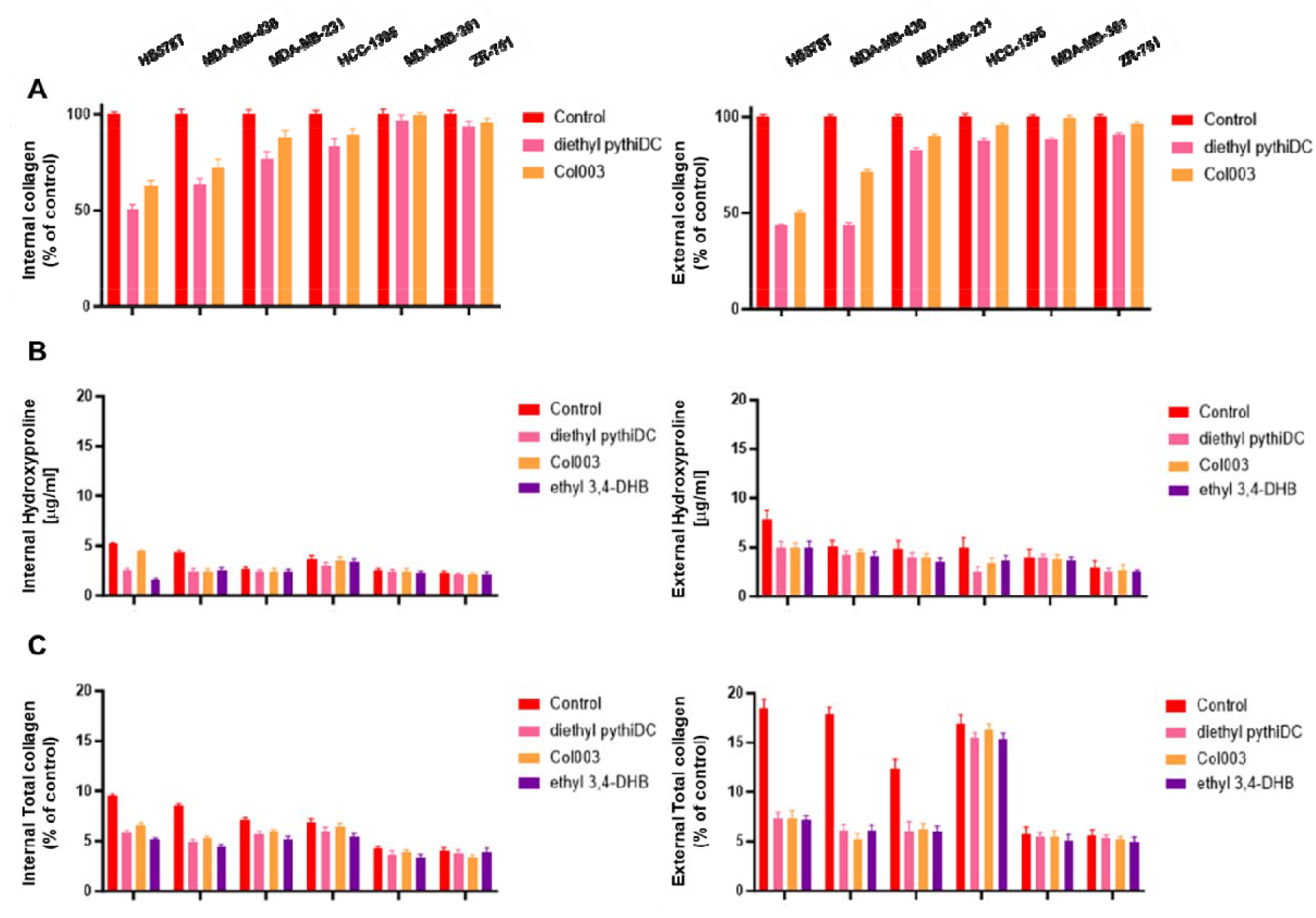
Collagen production. Quantification of internal and external secreted collagen was quantified without treatment (red) or with Col003 (orange), ethyl 3,4-DHB (purple) and diethyl pythiDC (pink) on IntClust-2 or non-IntClust-2 cells. **A**) COL1A1, **B**) Hydroxyproline and **C**) Sirius red detection were measured after 48 h of exposure to the drugs. The results are shown as mean ± standard deviation.

Accumulation of hydroxylated proline in collagen was diminished on some of the IntClust-2 cells but non-IntClust-2 were barely impacted by the P4HA3 and SERPINH1 drugs, either inside or outside the cell (**Fig. 5B**).

Levels of total collagen deposition were significantly reduced in the cells and the media of the IntClust-2 lines (**Fig. 5C**).

### P4HA3 pharmacologic inhibition effect on IntClust-2 xenograft tumor growth

The ‘acid test’ for oncoprotein inhibitors is to see if they are effective *in vivo*. As no adequate GEM models exist that mirror 11q13 amplification, we employed a xenograft approach.

To evaluate the role of P4HA3 in tumor growth we implanted subcutaneously IntClust-2 (HS578T) or non-IntClust-2 (ZR-751) cell lines into the right dorsal flanks of 6-week-old nude mice. Animals were observed for tumor growth, pain, and activity following all regulatory standards in accordance with the guidelines of the FCCC Institutional Animal Care and Use Committee. When the tumors were established (100 mm^3^) the therapy with a P4HA3 inhibitor (diethyl pythiDC) was initiated. Tumor-bearing mice were treated with vehicle or diethyl pythiDC (100 mg/kg/week) for a period of 4 weeks. Treatment with diethyl pythiDC significantly reduced the growth of HS578T xenografts (∼1500 mm^3^ versus ∼500 mm^3^), while mice with ZR-751 had a slight decrease of tumor growth (∼1050 mm^3^ versus ∼925 mm^3^) (**Fig. 6A**). The tumor volume was significantly smaller by about 70% when treating the HS578T mice compared to the control. The ZR-751 tumor volume had a mild effect (**Fig. 6B**). In both types of xenografts, the mice conserved the same body weight (**Fig. 6C**).

**Figure 6.**
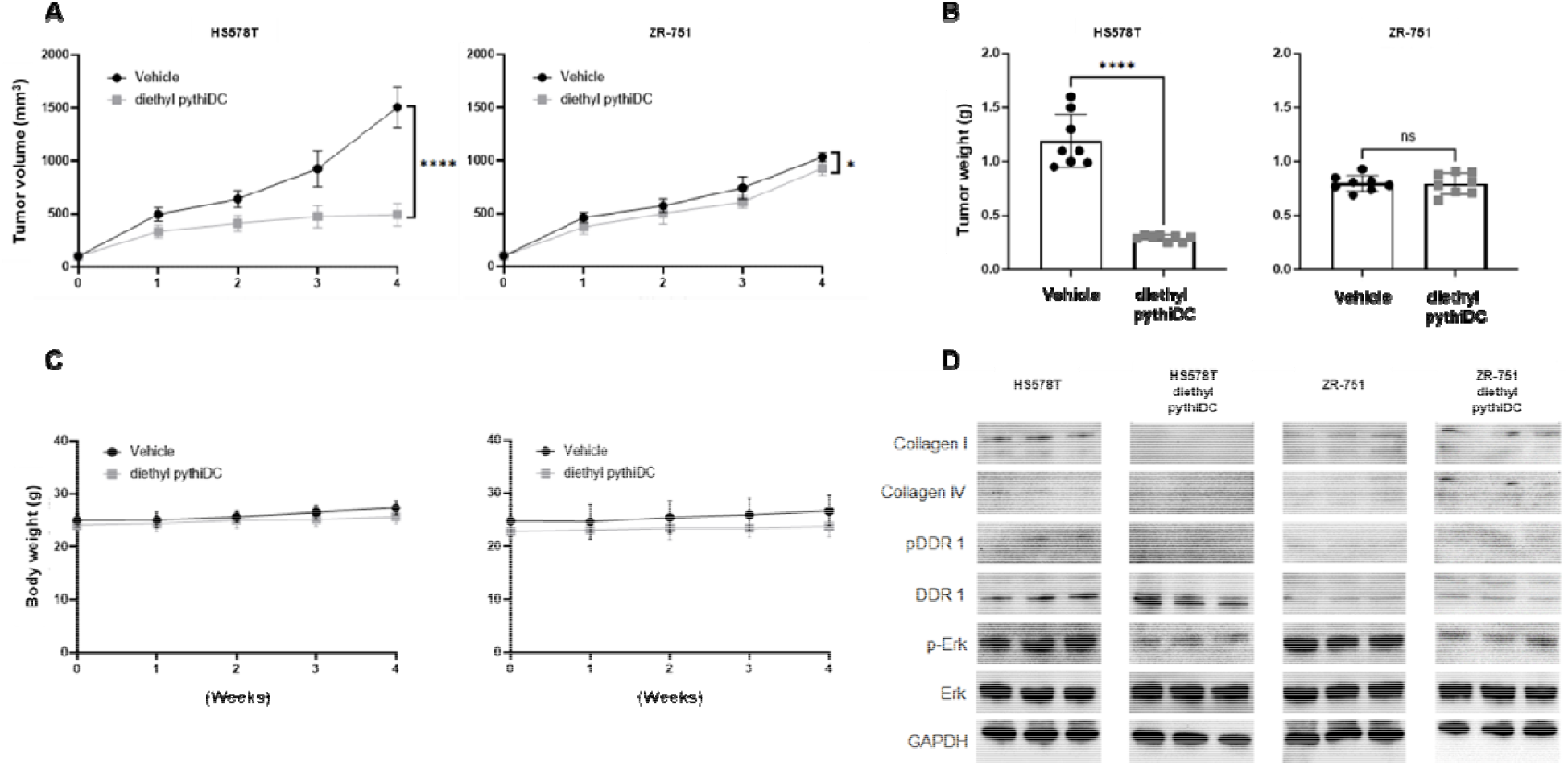
Inhibition of tumor growth *in vivo* by P4HA3 inhibitor. IntClust-2 (HS578T) and non-IntClust-2 cells were subcutaneously injected into the flanks of Nude mice (cohorts of eight mice/treatment). Ten days after inoculation, when the tumors reached a volume of at least ∼100 mm^3^, the animals were treated with vehicle (black) or diethyl pythiDC (gray) by IP for 4 weeks. **A**) Volumetric changes in tumor size between untreated mice and mice treated with inhibitor. **B**) Mice weight with vehicle and diethyl pythiDC and **C**) Tumor weight was measured after mice were sacrificed. **D)** Collagen I, collagen IV, pDDR1, DDR1, pErk and Erk activities were measured by Western blot for each condition and cell line. GAPDH was used as a loading control. Results are shown as the mean ± SD of at least three independent experiments (* p < 0.05, ** p < 0.01, *** p < 0.001, \*\*\*\**p* < 0.0001 versus control).

We determined the contribution of the P4HA3 inhibitor on the tumor’s related signaling pathways. The comparison between the treated mice of the two different xenografts was analyzed by WB. In the case of the HS578T tumors, we observed a reduction of collagen I, collagen IV, pDDR1 and p-ERK in the treated mice. The ZR-751 protein levels remained the same (**Fig. 6D**).

IHC was carried out to assess tumor cell morphology, collagen production, cell proliferation and death. HS578T tumors presented a significant loss of the proliferation marker (Ki-67) and the collagen production but an increase in the apoptotic (caspase-3) in the treated mice. No significant difference was observed between the control and treated ZR-751 animals (**Fig. 7A, B**).

**Figure 7.**
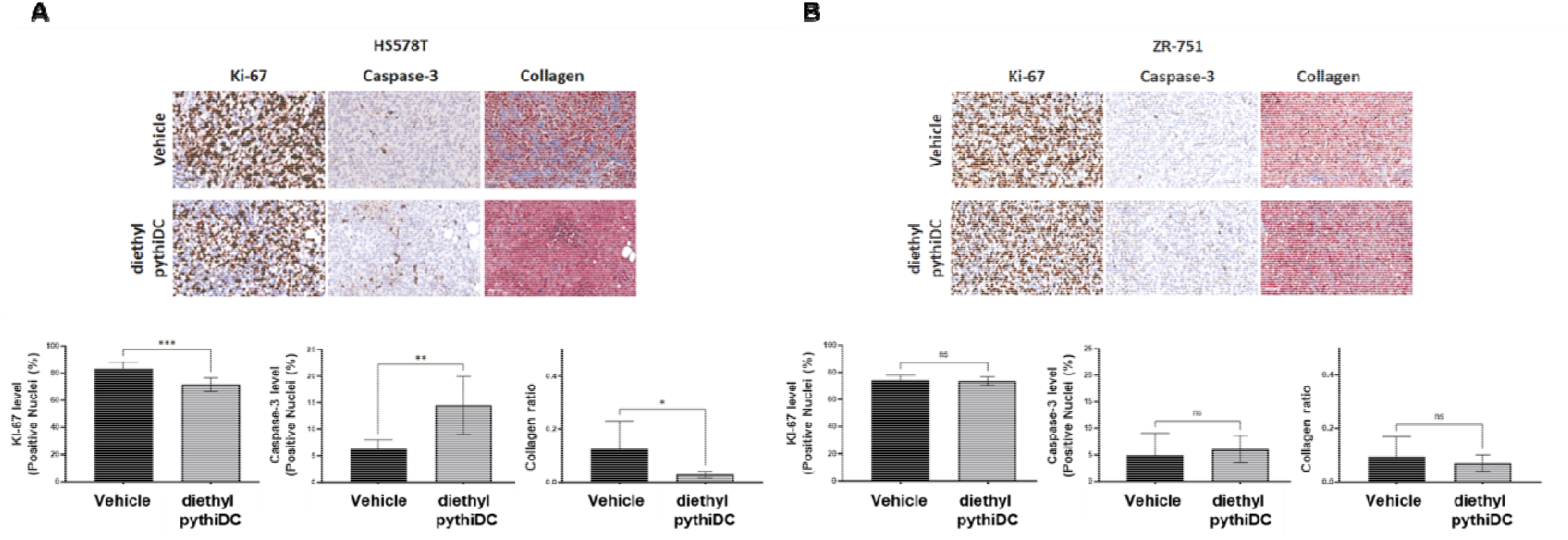
Representative tumor sections of IntClust-2 and non-IntClust-2 mice, either untreated or treated. Tumor tissues of **A)** IntClust-2 and **B**) non-IntClust-2 were embedded with paraffin; stained for markers of proliferation (Ki-67), apoptosis (cleaved caspase 3) and collagen production; and subjected to immunohistochemistry analysis (400×). Images and percentage quantification of intensity levels of caspase-3, Ki67 and collagen were determined by Student’s t-test. The results are shown as mean ± standard deviation (* p < 0.05, ** p < 0.01, *** p < 0.001, \*\*\*\**p* < 0.0001 versus control).

## Discussion

IntClust-2 tumors, affecting 3.1% of breast cancer patients, are highly resistance to current therapies and exhibit one of the worst prognoses among genomically-clustered groups, (1, 5, 9, 10). Consequently, these patients are a high priority for the development of novel therapeutic targets. Previous efforts to understand this type of breast cancer have mostly focused on well-known oncogenes such as *CCND1* and *PAK1* that reside within the defining 11q13/14 amplicon (5). However, it has become clear that neither of these genes is responsible for all the transforming activity within this region. While a few RNAi-based screens have been performed for genes in this region, these screens have focused on a few candidate drivers rather than on the amplicon as a whole.

To our knowledge, our work, using CRISPR to delete all genes within this region, is the first to systematically analyze the entire amplicon. Through this approach, we identified several previously ignored drivers, including *P4HA3* and *SERPINH1*, both encoding enzymes required for collagen processing (23). There are at least two potential limitations to our approach. First, recent reports suggest that using CRISPR to analyze amplified genes might be problematic, due to the generation of multiple double-stranded breaks in amplified DNA (18, 19). However, we were able to control these effects by rigorously comparing all targeted genes to each other, cancelling out any non-specific DNA cleavage effects. Second, it is probable that multiple driver genes within the 11q13/14 region contribute to oncogenesis. Our findings indicate that, in addition to the well-recognized driver functions of *CCND1* and *PAK1*, both *P4HA3* and *SERPINH1* represent a new and heretofore unrecognized set of oncogenic drivers in the 11q13/14 amplicon. As this chromosomal region is also frequently amplified in ovarian (16) cancer and head and neck cancer (24) findings may have broad implications in several malignancies.

Clinical observations have shown that breast cancer is often characterized by a dense reactive stroma associated with extensive collagen I deposits (25) and that alterations in the organization of collagen I occur during the initial stages of breast cancer development and promote local invasion (26). There are different genes and therefore proteins required for collagen synthesis. Along these lines, it is interesting to note that expression of P4HA1 and P4HA2 has been reported to be essential for breast cancer metastases (27), promoting cancer cell alignment along collagen fibers and resulting in enhanced invasion and metastasis to lymph nodes and lungs (28, 29). In accord with these ideas, treatment with ethyl 3,4-dihydroxybenzoate, a P4HA inhibitor, has been shown to decrease tumor fibrosis and metastasis in a mouse model of breast cancer (23, 28, 30). Similar observations have been made using compounds that inhibit P4HA activity by blocking α-ketoglutarate production (31).

Knowing that collagen deposition as well as the amplification of the 11q13.2-14.1 chromosomal region (IntClust-2) due to combined overexpression of several oncogenes in this amplicon are responsible for the poor outcome of breast cancer patients, we decided to study by gene-editing if there were potential oncogenes within the cluster associated to collagen production and signaling. Our aim was to assess if our findings apply to IntClust-2 tumors in general, and if genetic and/or pharmacologic manipulation of collagen synthesis and/or signaling would provide new targets for anti-cancer therapy in this challenging, difficult-to-treat, poor prognosis form of breast cancer.

We demonstrate that tumor cells bearing 11q13/14 amplifications generally express increased amounts of SERPINH1, P4HA3, and collagen, as well as exhibiting altered matrices and enhanced DDR signaling. Having established the basal signaling state of the IntClust-2 cell lines, we probed the two main arms of collagen signal transduction to determine whether either or both are required to support the proliferation, survival, and/or motility of these cells. We found that a panel of IntClust-2 cells were sensitive to knockdown of either P4HA3 or SERPINH1, respectively. Additionally, we explored whether this heightened sensitivity persists when using small molecule inhibitors instead of siRNA.

Gilkes *et al*. previously reported that a prolyl hydroxylase inhibitor, ethyl 3,4-DHB, inhibited collagen secretion by the IntClust-2 cell line MDA-MB-231 (28). We evaluated the effects of this compound, along with another P4HA inhibitor, the esterified biheteroaryl dicarboxylate (diethyl pythiDC) (23), on collagen synthesis, cell proliferation, survival, and motility. As a parallel approach, we targeted the collagen chaperone SERPINH1/Hsp47 using the small molecule inhibitor Col003 (32). In both cases, we observed that all the compounds specifically impacted oncogenic processes of IntClust-2 cell lines.

Our model suggests that IntClust-2 breast cancer cells over-produce and are over-stimulated by collagen related proteins. This feature presents a therapeutic opportunity. Previous inhibitors of collagen prolyl hydroxylases had been studied against different types of cancer but presented undesirable effects due to iron chelation, affecting multiple cellular processes (23). We therefore used diethyl pythiDC *in vivo*, a compound that selectively targets collagen prolyl 4-hydroxylases without affecting other prolyl hydroxylases such as HIFα or Jumanji proteins (23, 29). This molecule had a clearly beneficial impact in an IntClust-2 xenografts model without undue toxicity.

These studies demonstrate that collagen deposition and tissue stiffness are negative prognostic signs in breast cancer. Therefore, we propose a molecular pathway that is potentially druggable in 11q13/14 amplified tumors, by impeding collagen prolyl hydroxylases such as P4HA3 or collagen chaperones such as SerpinH1/Hsp47. In addition, it may be possible to block the collagen pathway further downstream using small molecule inhibitors of integrin function or for the collagen-stimulated kinase DDR1/2.

## Authors’ Contributions

D. Araiza-Olivera: Conceptualization, data curation, formal analysis, investigation, validation, methodology, visualization, writing–original draft, writing– review and editing. T.Y Prudnikova: Conceptualization, data curation, formal analysis, validation, investigation, visualization, methodology. C. Uribe-Alvarez: Data curation, formal analysis, validation, investigation, visualization, methodology. K.O. Cai: Data curation, formal analysis, validation, investigation, visualization, methodology. R.T. Raines: provision of diethyl pythiDC, writing– review and editing. J. Franco-Barraza: Data curation, formal analysis, validation, investigation, visualization, methodology. J. Chernoff: Conceptualization, resources, supervision, funding acquisition, validation, investigation, visualization, methodology, writing–original draft, project administration, writing–review and editing.

## Acknowledgments

We thank Dr. Neil Johnson for cell lines (HCC1395 and MDA-MB-231). We would also like to thank Fox Chase Cancer Center core facilities including histopathology, laboratory animal and cell culture facilities.

This work was supported by grants from the PA Breast Cancer Coalition (JC), R35 GM148220 (RTR), and NCI Core Grant P30 CA06927 (Fox Chase Cancer Center).

