## Supplementary data for "Identifying and targeting key driver genes for collagen production within the 11q13/14 breast cancer amplicon"

**Supplemental information**

**Supplemental Table 1. sgRNA sequences and drop-out screen results.**

```
gRNA_ID	Gene_Symbol		Sequence			sum52/cas9 	sum52/cas9
day 0		day 14
CCND1_1	CCND1			GGTTGGCATCGGGGTACGCG	27437		12598
CCND1_2	CCND1			GTTCGTGGCCTCTAAGATGA	55973		25846
CCND1_3	CCND1			GATGTCCACGTCCCGCACGT	74430		14247
CCND1_4	CCND1			GTCGTTGAGGAGGTTGGCAT	51719		22048
CCND1_5	CCND1			GTTCGTGGCCTCTAAGATGA	55973		25846
CCND1_6	CCND1			GATGTCCACGTCCCGCACGT	74430		4247
ORAOV1_1	ORAOV1		GCACAGTTGCACCACTGAGA	42319		75936
ORAOV1_2	ORAOV1		GGCGGATGAGAGGTTTCATG	9830		23902
ORAOV1_3	ORAOV1		GGTTTCATGGGGAAGGGTAT	29426		31125
ORAOV1_4	ORAOV1		GGTCATCATAAGGGAATTTC	48581		131586
ORAOV1_5	ORAOV1		GTTGCACCACTGAGAAGGAC	25528		82302
ORAOV1_6	ORAOV1		GAAGACTTAGACAAGATCAG	62575		72096
FGF19_1	FGF19			GCGCCGACGGCAAGATGCAG	33886		22802
FGF19_2	FGF19			GGTGGTCCACGTATGGATCC	24387		37042
FGF19_3	FGF19			GTGTCTTGGGCACCTACCCG	21103		12079
FGF19_4	FGF19			GTACAGGTGCCGCAGGCGGA	62156		121819
FGF19_5	FGF19			GATCCTGGCCGGCCTCTGGC	38131		19839
FGF19_6	FGF19			GAAGCAGCTGGAGAGCCCGT	101466	22901
FGF4_1	FGF4			GTCGCCGGCGCCGCTCTGGA	38666		61742
FGF4_2	FGF4			GCGCCAACGAGAGCGCCACC	38084		12754
FGF4_3	FGF4			GCAAGGCCAGCAGGACCGCC	43320		99018
FGF4_4	FGF4			GGCCGAGCTGGAGCGCCGCT	26000		30965
FGF4_5	FGF4			GCGCGGTTGCGGCGCGATGC	42669		1685
FGF3_1	FGF3			GAGGGGACGACTCTATGCTT	1270		23834
FGF3_2	FGF3			GCTATAATACGTATGCCTCC	17403		97041
FGF3_3	FGF3			GGCGGCGTCTACGAGCACCT	19129		62136
FGF3_4	FGF3			GCAGTAGAGCTTGCGGCGCC 	0		0
FGF3_5	FGF3			GGCGCGGTTGCGGCGCGATG	42993		153910
FGF3_6	FGF3			GCACTCACCGTGCCCTCGGA	45877		48150
TMEM16A_1	TMEM16A		GGCCACGGGTCTCATTAATG	77667		34133
TMEM16A_2	TMEM16A		GCCCCCGATGGACTACCACG	15272		33400
TMEM16A_3	TMEM16A		GTACTCGACGCTCCCGGCCG	34512		45101
TMEM16A_4	TMEM16A		GCCCGCACTCACCGTGCCCT	49789		148912
TMEM16A_5	TMEM16A		GATGTGGACGCTGCGGTCCT	30124		49227
TMEM16A_6	TMEM16A		GAGCTCAAGTTCCTATGCCT	35398		33242
FADD_1	FADD			GAGCAGCTCGCGCAGGAGCT	34472		19836
FADD_2	FADD			GCGGCGCGTCGACGACTTCG	49406		33548
FADD_3	FADD			GTTCCTGGTGCTGCTGCACT	13215		39915
FADD_4	FADD			GCGCGTGGGCAAGCGCAAGC	36810		12637
FADD_5	FADD			GTCGTTCTGCTCCAGCAGCA 	0		0
FADD_6	FADD			GCTTCACTTCATCATGGGTA	34438		26439
PPF1A1_1	PPF1A1		GAGAGAGACTCAAGAAACGC	54729		65928
PPF1A1_2	PPF1A1		GGTCTCCATGCTAGAAGAAA	58791		51399
PPF1A1_3	PPF1A1		GAAACGCTGGCCTTAACCCA	42024		44409
PPF1A1_4	PPF1A1		gAGAGGTGCTTCACTTCATCA	0		0
PPF1A1_5	PPF1A1		GCCAAGACGGTGCAGGGCTC 	0		0
CTTN_1	CTTN			GCAAAGATGGGGTGCCAAGA	31971		57501
CTTN_2	CTTN			GTCCACCTGCGAGCAGTGCT	19099		39919
CTTN_3	CTTN			GTCCATCGCCCAGGATGACG	11873		13847
CTTN_4	CTTN			GGTGCAGGGCTCCGGGCACC 	0		0
CTTN_5	CTTN			GCATCAGACCCTTAAGGAGA	50879		52863
CTTN_6	CTTN			GGCAAGATCCTTACCAGGCA	47186		132987
DHCR7_1	DHCR7			GAAGTAGTAGACGATGAAGG	38382		61952
DHCR7_2	DHCR7			GAACCGCATCTCAAGGGCAG	46244		6095
DHCR7_3	DHCR7			GCCATCTAGACTCTTGGCTT	41408		31419
DHCR7_4	DHCR7			GATGAAGTAGTAGACGATGA	19775		6000
DHCR7_5	DHCR7			GTCCGAGAGCCGAGCATGTC	61320		84066
DHCR7_6	DHCR7			GCAAATTGCCCTCGAAGTCC	12285		95551
NADSYN1_1	NADSYN1		GCAGAGTGATATTCCTCAAC	110678	72564
NADSYN1_2	NADSYN1		GCACTCGTTTCAAGTCCTAG	42208		58293
NADSYN1_3	NADSYN1		GATGGGCCGGAAGGTGACCG	22362		3875
NADSYN1_4	NADSYN1		GAGACTCCACAAGGGCCGCT	2659		8500
NADSYN1_5	NADSYN1		GAGTGACGGGAGACTCCACA	5198		48260
NADSYN1_6	NADSYN1		GCAGAGTGATATTCCTCAAC	110678	72564
KRTAP5-7_1	KRTAP5-7		GGGGCTGTGGCTCTGGCTGT	8952		14913
KRTAP5-7_2	KRTAP5-7		GGCTGCTGTGGCTGTTCCGA	75136		36112
KRTAP5-7_3	KRTAP5-7		AAGGCTGTGGCTCCGGCTGT	17638		14246
KRTAP5-9_1	KRTAP5-9		GCACACAGGGACACAGCATC	15911		95121
KRTAP5-9_2	KRTAP5-9		GCAGGATGACCCACGACCCG	43616		116463
KRTAP5-9_3	KRTAP5-9		GTTGCTCCTCCTCGGGTCGT	25194		143809
KRTAP5-9_4	KRTAP5-9		GTAGACGGGTGCACAGCAGC	56196		119654
KRTAP5-9_5	KRTAP5-9		GGATGACCCACGACCCGAGG	52644		111831
KRTAP5-9_6	KRTAP5-9		AAGGTGGCTGTGGTTCCTGT	61468		48883
KRTAP5-10_1	KRTAP5-10		GCAGCAGGATCCACAGCCTG	45672		8720
KRTAP5-10_2	KRTAP5-10		GAGGCTGTGGCTCCGGCTGT	47744		9443
KRTAP5-10_3	KRTAP5-10		ACAGGAACCACAGCCACCTT	39190		32103
KRTAP5-10_4	KRTAP5-10		GGGCTTACAGCAATTGCACT	36995		2300
KRTAP5-11_1	KRTAP5-11		GCAGCAGGGTTTGGAGCAGC	9068		48865
KRTAP5-11_2	KRTAP5-11		GCCAGAATCATGGGCTGCTG	63062		32181
KRTAP5-11_3	KRTAP5-11		GGCTGTGGCTCCGGCAGTGG	5646		8966
KRTAP5-11_4	KRTAP5-11		GGGCTTACAGCAGTTGGACT	16595		28647
KRTAP5-11_5	KRTAP5-11		GAATGACCCACAGCCTGAGG	16660		67452
KRTAP5-11_6	KRTAP5-11		GGATTGGCAGCAGCAGGGTT	59620		77263
FLJ10661_1	FLJ10661		GCGGGCTCCTCAATTCTACA	44973		71985
FLJ10661_2	FLJ10661		GTAAGGGCCATGTAGAATTG	35641		14476
FLJ10661_3	FLJ10661		GCGCCCTGCGCTCCTTCCGC	43771		85483
DEFB108B_1	DEFB108B		GAGATCTGTGAACGTCCAAA	26077		89350
DEFB108B_2	DEFB108B		CTTGAAACAGAAATCCATGT	31603		77774
RNF121_1	RNF121		GGATCATTGAGAACACGTAT	10903		19335
RNF121_2	RNF121		GTTCCAGAACTCCATAGTAG	41372		78032
RNF121_3	RNF121		GGCAGGATACACATGATTGC	16839		63789
IL18BP_1	IL18BP		GCTGTGGTGGTCTGCGAGAC	8488		108641
IL18BP_2	IL18BP		GACCAGGAGAGTGACGACGT	9435		33661
IL18BP_3	IL18BP		GGAAGTGACCTGGCCAGAGG	14746		38914
IL18BP_4	IL18BP		GTCCTTTGTGCTTCTAACTG	57893		90165
IL18BP_5	IL18BP		GCTTAGCTGCTGGGAACACT	21127		74675
IL18BP_6	IL18BP		GTCCAGCATTGGAAGTGACC	31080		21572
NUMA1_1	NUMA1			GCTGTCCCTCTTCAGTGCCA	65158		45609
NUMA1_2	NUMA1			GTCTGCAGGATATCACCCAT	49836		10119
NUMA1_3	NUMA1			GGAGGCTGTGCTGCAGCTCC	38589		78085
NUMA1_4	NUMA1			GAGATGCTGGAGACTGAGCG	32402		335221
NUMA1_5	NUMA1			GGTTGGAGATGGAGCTCTGC	65560		108130
NUMA1_6	NUMA1			GGTGAGCTCAGAGGTCAGAG	25248		62452
LRRC51_1	LRRC51		GGGGCTGACCTTGAATGACG	20759		10660
LRRC51_2	LRRC51		GACCCCACTGAAGTCGAACG	4669		37077
FLJ20625_1	FLJ20625		GTAGGCATAAGCAGCTATCC	12027		40576
FLJ20625_2	FLJ20625		GGCCTGCTCATCAGTGCGAG	62647		40006
DKFZP564M082_1/DKFZP564M082	GAGCCACTCACCTCTGAGGC	8764		1570
DKFZP564M082_2/DKFZP564M082	GGACATGGAAGGCAACGAAC	21981		7984
FOLR3_1	FOLR3			GGGCTGTGTCTTGTGGTGCT	47604		25961
FOLR3_2	FOLR3			GCACTCAGAGACAGCTGTCC	8610		42283
FOLR3_3	FOLR3			GAATCAATAATCCCACGAGA	61329		40709
FOLR3_4	FOLR3			GACATTGAGCAGGTCCGTCC	28186		32229
FOLR3_5	FOLR3			GACAGCTGTCCTGGATAAAG	5659		41778
FOLR3_6	FOLR3			GGAACAGCACTCATACCTGC	0		0
FOLR2_1	FOLR2			GCCAGTTGCTCTTGCACGTG	43475		12358
FOLR2_2	FOLR2			GGATGTGCCCTTATGCAAAG	34752		31490
FOLR2_3	FOLR2			GAACCTCGCCACTTCCTCGT	14661		10854
FOLR2_4	FOLR2			GGTCCCAGTTAAAGTTGTAC	0		0
FOLR2_5	FOLR2			GGGCCCTGGATCCAGCAGGT	0		1416
FOLR2_6	FOLR2			GCAAAGAGGACTGTCAGCGC	9330		7504
INPPL1_1	INPPL1		GATGGTCTTGGCCTTACGTG	94019		65709
INPPL1_2	INPPL1		GGTCCGAGACAGCGAGAGCG	20413		29724
INPPL1_3	INPPL1		GTACCACCGCGACCTGAGCC	17465		6944
INPPL1_4	INPPL1		GCCTGATGGAGAAGATTTCT	0		1450
INPPL1_5	INPPL1		GTGCGCCGCTTCCAGACCCT	5016		33958
INPPL1_6	INPPL1		GGCGCACACAAGGCCCTGGT	10694		2
PHOX2A_1	PHOX2A		GTACAGGTCCTGCTTGCTGC	25742		6416
PHOX2A_2	PHOX2A		GCCATGGAGGCGTCCGCCTA	44379		58256
PHOX2A_3	PHOX2A		GCCGTAGGCGGACGCCTCCA	39625		1215
PHOX2A_4	PHOX2A		GGGCTGTATTGGAAGCCGCC	0		9043
PHOX2A_5	PHOX2A		GCAGTTGGAGGAGCCGAGCG	87727		133198
PHOX2A_6	PHOX2A		GCTGCGCTCACCTGCCGAGT	33703		89451
SKD3_1	SKD3			GGCTGAGGGTAGCCACCGGG	5399		7888
SKD3_2	SKD3			GGCTACTCCTCCGGCTGCTC	21934		6753
SKD3_3	SKD3			GGCCTTGTTCTCCGGACGTG	20607		5726
SKD3_4	SKD3			GCAATGAACCACCAGCGCTG	1		1029
SKD3_5	SKD3			GTGTTTCTTCGGGACCAGGA	1		1
SKD3_6	SKD3			GATCTCATAATCACCCTTGT	0		15003
PDE2A_1	PDE2A			GAAACTCAAGTGTGAGTGCC	28877		14420
PDE2A_2	PDE2A			GAGGATCTTGCGGTCGCGGT	27706		29037
PDE2A_3	PDE2A			GCTGCCACTGGGAGACAGGA	52122		5939
PDE2A_4	PDE2A			GGCTCTGTCATCGACATTTC	1		0
PDE2A_5	PDE2A			GTCTACACCTACCTACTGGA	0		4427
PDE2A_6	PDE2A			GCGGCTGGGCTGCAATGGGC	7175		4584
CENTD2_1	CENTD2		GTAGCATGCCCATGTCCATC	50820		44981
CENTD2_2	CENTD2		GAATAAGCTGTACGTGGCCG	31609		88860
CENTD2_3	CENTD2		GTGGCTGCGGGCATTGCACC	16767		33980
CENTD2_4	CENTD2		GCCCGTGGGACTCGGCTCGG	0		0
CENTD2_5	CENTD2		GTGGCCAGTCTGCTGAGCGA	26956		2915
CENTD2_6	CENTD2		GAGGATGACCACGCCTATGA 	0		0
STARD10_1	STARD10		GCTCGGACCAAGTCTTTCCG	29535		5531
STARD10_2	STARD10		GTCTGTCTGGGTGCAGGCTG	5157		9004
STARD10_3	STARD10		GCGAAAGTCTTGGTCATCGG	10066		16795
STARD10_4	STARD10		GCTGGAACCTGACCTATAGC	10421		2
STARD10_5	STARD10		GTCAGCGTTGACTGTCAAGC	64519		23064
STARD10_6	STARD10		GATGACATCACGGTTCTTCA	0		1
FLJ00012_1	FLJ00012		GCTCCGACTCCAGCGTGCCC	4017		5627
FLJ00012_2	FLJ00012		GACATACCTGAACACAGCTT	10010		22055
FLJ00012_3	FLJ00012		GCTGGAAACGCCACATCGTG	46119		63133
FCHSD2_1	FCHSD2		GTCAATACCTGAAGAGAGAT	49799		58774
FCHSD2_2	FCHSD2		GGAATTACACTCAGATCGCC	5641		56128
FCHSD2_3	FCHSD2		GCCGCCGCCGAGGAAGGTAA	17018		12800
FCHSD2_4	FCHSD2		GTGATTTGCTTGAAGATATG	0		22455
FCHSD2_5	FCHSD2		GAAGATAATCATTCCTTGCG	10174		2963
FCHSD2_6	FCHSD2		GATGTGCATTTGCTGCCGCT	2870		130412
P2RY2_1	P2RY2			GCTGCCTGTGTCCTACGGCG	22053		56835
P2RY2_2	P2RY2			GGTGCGCACGGACTTGCGCT	19307		3301
P2RY2_3	P2RY2			GTCCGTGCGCACCATCGCCG	17771		61738
P2RY2_4	P2RY2			GTAATAGACCAGCAGCGGCA	0		0
P2RY2_5	P2RY2			GAAGGGCCAGTGGTCGCCGC	0		3
P2RY2_6	P2RY2			GAGGATGCTGCAGTAAAGGT	10386		1
P2RY6_1	P2RY6			GCCAGCACCGCCGAATACAC	19435		3149
P2RY6_2	P2RY6			GTTCTCGCGGTAGACACAGG	22169		48976
P2RY6_3	P2RY6			GAAGTTCTCGCGGTAGACAC	36798		23165
P2RY6_4	P2RY6			GTAATGACACAGATGTTCAG	60807		32953
P2RY6_5	P2RY6			GTACACGGCCGTGCGGGTCA	49487		35206
P2RY6_6	P2RY6			GTACACCCTAAACCTTGCTC	40618		8280
ARHGEF17_1	ARHGEF17		GGACACAGAGCAGTCGTATG	37441		24210
ARHGEF17_2	ARHGEF17		GCAGCTTGAACGAGACGCTG	33849		12936
ARHGEF17_3	ARHGEF17		GTTACGCCACTATGGCGGAC	4		5664
ARHGEF17_4	ARHGEF17		gTTCACCAGAAGCTCGTAGCG	0		0
ARHGEF17_5	ARHGEF17		gCACTCACCGACTGGACGAGC	0		0
ARHGEF17_6	ARHGEF17		gAAGATCTCGTCCACCAGTGA	0		0
TNFRSF19L_1	TNFRSF19L		GGCGATGACCGCGTACTGGG	49761		19758
TNFRSF19L_2	TNFRSF19L		GCCAAAGGGTTGTTGATGTC	19877		8955
TNFRSF19L_3	TNFRSF19L		GCAACTCGAGATACACTCTG	15257		22947
TNFRSF19L_4	TNFRSF19L		GTCAGGGTGGCGAGAGGCCA	19937		2
TNFRSF19L_5	TNFRSF19L		GGCCCGACGTGGCGTGGAGG	0		1
TNFRSF19L_6	TNFRSF19L		GGCAGCCTGGGAACGGCACC	4691		3823
KIAA0280_1	KIAA0280		GTATTTACTGAAAGCCGCAG	15642		21404
KIAA0280_2	KIAA0280		GTTTACAGCCCCGTGCAGCC	21270		2218
KIAA0280_3	KIAA0280		GTAAGTTCGGTTCTCGGTCC	39389		79908
KIAA0280_4	KIAA0280		GCATAACTGGGACTATTGGT	0		0
KIAA0280_5	KIAA0280		GGAAGACGAGTTCTGTGGCC	0		0
KIAA0280_6	KIAA0280		GAAGGTGCCTTCAGTGCCAC	18961		101419
PLEKHB1_1	PLEKHB1		GCCTGCTGACTGTGAACCTA	14340		43012
PLEKHB1_2	PLEKHB1		GACTCACAGGGCATCATCCT	29798		67070
PLEKHB1_3	PLEKHB1		GAACTGGTTTGCCCTGTGGC	25516		30097
PLEKHB1_4	PLEKHB1		GGTTTGCCCTGTGGCTGGAC	0		0
PLEKHB1_5	PLEKHB1		GGGATTTGCTTTGCAGAGCA	7644		4173
PLEKHB1_6	PLEKHB1		GCACCTCTGTGCGGAGACCA	42674		3619
RAB6A_1	RAB6A			GCTACACGTCGAAAGAGCTG	20812		18668
RAB6A_2	RAB6A			TCACTGACTGGTTGCTCCTG	1		0
MRPL48_1	MRPL48		GGAAGTGAGAGCCATTAATT	72629		54037
MRPL48_2	MRPL48		GCTTGTACTTTCCAATGCCG	32854		17522
MRPL48_3	MRPL48		GAATTCTACTAAGTATCAGT	23450		42679
MRPL48_4	MRPL48		GTACTTTCCAATGCCGTGGG	20389		25658
MRPL48_5	MRPL48		GCACTTAATTAAAGCAGAAG	1		1749
MRPL48_6	MRPL48		GACTGCATATGATATGACCC	20542		1444
E2IG2_1	E2IG2			GACCAGCTGATCTCCCGCTC	21451		16187
E2IG2_2	E2IG2			GGGTCCAGGTATGGCCTTGA	47486		14678
E2IG2_3	E2IG2			GCACTCCTGCACTGCAAAGT	83348		13332
E2IG2_4	E2IG2			GCTGGCATTGCCGCCAGTCC	27164		4698
E2IG2_5	E2IG2			GGTGAAGAAAGACGATGAGG	1		9549
E2IG2_6	E2IG2			GGTATGGCCTTGAGGGACTG	5669		41219
WDR71_1	WDR71			GTACAAGTCATCCACAGATC	11123		39456
WDR71_2	WDR71			GATCAACGATGGCTGTATCC	22197		61795
WDR71_3	WDR71			GGATACAGCCATCGTTGATC	15572		30985
WDR71_4	WDR71			GGCTTCCAATGGAGAACTCA	63768		51634
WDR71_5	WDR71			GTGACCTTCAAAGGTCACAA	37465		5787
WDR71_6	WDR71			GTGGTGTCTGCTTCTCGAGA	24913		17380
DNAJB13_1	DNAJB13		GGTAGAGATCCCGTTCGACT	15333		34560
DNAJB13_2	DNAJB13		GTAGAGATCCCGTTCGACTT	25042		40536
DNAJB13_3	DNAJB13		GGTGCTTAAGGGCGAGTCTG	36289		36065
DNAJB13_4	DNAJB13		GTGTTCCACGAGTTCTTTGG	38488		19922
DNAJB13_5	DNAJB13		GTCGTAGATGCCTCTCTTCA	3861		5748
DNAJB13_6	DNAJB13		GGTGGGATTCCTTTGGAGTT	27199		52624
UCP2_1	UCP2			GCCCAGTACCGCGGTGTGAT	61687		58665
UCP2_2	UCP2			GCGCTGGCTGTAGCGCGCAC	10881		1
UCP2_3	UCP2			GGTGATGAGATCTGCGATGC	20391		53884
UCP2_4	UCP2			GAAACTTCACAGTGGCAGTA	20098		43226
UCP2_5	UCP2			GCAGATCCAAGGAGAAAGTC	12240		6139
UCP2_6	UCP2			GCCCATCACACCGCGGTACT	1		34764
UCP3_1	UCP3			GCCGGGCCGTCTGGACCGCC	38206		41308
UCP3_2	UCP3			GGAGAACCAGGCGGTCCAGA	30534		13654
UCP3_3	UCP3			GCCACGGTACTGCACGAGCC	36102		16752
UCP3_4	UCP3			GCCAAGGTCCGCCTGCAGGT	10932		4850
UCP3_5	UCP3			GTGCAGTACCGTGGCGTGCT	21876		3425
UCP3_6	UCP3			GCCGGCCACCAGCCCATTGT	13826		8629
DKFZP586P0123_1/DKFZP586P0123	GCAGTAGATCAACCACTCCG	58767		25909
DKFZP586P0123_2/DKFZP586P0123	GTGTACTTGTCCGAGTGAGA	5826		97394
DKFZP586P0123_3/DKFZP586P0123	GTATGGTAATTACCTCGGAG	66011		15491
PME-1_1	PME-1			GAGCGGAGCCAAGATGCGAA	11292		90407
PME-1_2	PME-1			GAGGTCATTCTGCCCTTTCT	14136		61525
PME-1_3	PME-1			GTTTCTGCAGACAGATCTTC	6376		86918
PME-1_4	PME-1			GCCCACCTCTACCCGGCAGC	14323		1704
PME-1_5	PME-1			GTCTACAAGAGTGGTTCAGA	16935		2
PME-1_6	PME-1			GCTTTGGATCTGCGAAGTCA	0		9125
P4HA3_1	P4HA3			GCGGTGCTGGCGCTCGGGAC	8839		11160
P4HA3_2	P4HA3			GAGGCTGTCAGTCTCTTCCG	26776		91683
P4HA3_3	P4HA3			GAGGAATGTGGTACATAGTC	20077		17113
P4HA3_4	P4HA3			GAATGTGGTACATAGTCTGG	27946		732
P4HA3_5	P4HA3			GGCAGCAATGCGGTGGTTGA	24377		1607
P4HA3_6	P4HA3			GTCAACAGTGTCCTTCAGCC	21039		409
PGM2L1_1	PGM2L1		GTCTTCCTTGCGGTTGTGAG	46893		27314
PGM2L1_2	PGM2L1		GTTACGGAATGGGATGAACA	25015		86511
PGM2L1_3	PGM2L1		GCAGGACTTCGTTCTGCCAT	5945		12817
PGM2L1_4	PGM2L1		GCAGTGGCTCCGCTGGGATA	5403		6210
PGM2L1_5	PGM2L1		GTGTTGACTGTATTACTGTA	53416		18396
PGM2L1_6	PGM2L1		GAATGTGTGGAACCCTGGAA	22111		9076
KCNE3_1	KCNE3			GAGCCTTCAGCACGGCATGC	60137		155539
KCNE3_2	KCNE3			GGCATGCAGGCTCTCATACC	23407		110353
KCNE3_3	KCNE3			GCAATTTGCTCTGCCGGCCA	0		0
KCNE3_4	KCNE3			GAGAGGCGGGCCAGCCTACC	0		7845
KCNE3_5	KCNE3			GCTTGTCCACTTTGCGGGAG	5043		5390
POLD3_1	POLD3			GCTGGCAGTCACAGCTAGCT	12267		1267
CHRDL2_1	CHRDL2		GCAGAACCTCACACTCCCTC	35516		7509
CHRDL2_2	CHRDL2		GATACTCCCCCGGCGAGAGC	12464		40705
CHRDL2_3	CHRDL2		GAGGCGGTAACAACTCACAT	29513		1292
CHRDL2_4	CHRDL2		GATCTCTCCGTGTTGGTACA	2		0
CHRDL2_5	CHRDL2		GCTGCAGAGGACACACTGGT	21868		9637
CHRDL2_6	CHRDL2		GGGTGCTGGGCAGCCTGGTT	0		1969
KIAA1991_1	KIAA1991		GGCTGCAGGTCCGAGTACTC	12672		53409
KIAA1991_2	KIAA1991		GCGGAAGAGGCCCGAGTACT	55048		64278
KIAA1991_3	KIAA1991		GAGTACTCGGGCCTCTTCCG	46687		75409
XRRA1_1	XRRA1			GACTTGGTTGTCGTCATGCG	45919		32986
XRRA1_2	XRRA1			GTTGCTTTGCCAGCCTGGCT	66677		88435
XRRA1_3	XRRA1			GACGAGTCAGTAGACTGGAA	5520		31002
XRRA1_4	XRRA1			GAGCTTCCTCCAGGAGCGAC	34087		11001
XRRA1_5	XRRA1			GCTTGGACTGCAGCATGAAT	5596		2213
XRRA1_6	XRRA1			GTCCAAGCCAAGGATGCTTG	27486		21206
SPCS2_1	SPCS2			GCGGCGGCAGCTGTACAGGG	32323		8645
SPCS2_2	SPCS2			GACAGCTGCCAAATATCATC	4461		8244
SPCS2_3	SPCS2			GACCATTTATACCTCATATA	31485		22593
SPCS2_4	SPCS2			GCTGTACAGGGCGGGAGAAG	0		10989
SPCS2_5	SPCS2			GCCACTTCCTGTCCCGCAGT	9623		25075
SPCS2_6	SPCS2			GCGGCTTGTTGGATAAGGTG	1		1743
NEU3_1	NEU3			GGATGTAGTAGGTATACGCA	14488		1674
NEU3_2	NEU3			GCGCCCCATGGAAGAATCCC	26131		11491
NEU3_3	NEU3			GAGCGTCAACAGATTGTGTC	47633		20728
NEU3_4	NEU3			GAGCAGGGCTGGGATCCGGT	27046		26906
NEU3_5	NEU3			GCGTTCTACGAGGAGAGATG	20267		36912
NEU3_6	NEU3			GGCTTCCATCAGTGGCTTCA	25146		21039
OR2AT4_1	OR2AT4		GATGGGTAATGCCCTGATCC	26185		64806
OR2AT4_2	OR2AT4		GGAACTGATGCGAAGCACTG	12269		5959
OR2AT4_3	OR2AT4		GCCCATGATATGGAAGTCAA	28397		23046
OR2AT4_4	OR2AT4		GGATCAGGGCATTACCCATC	23457		33724
OR2AT4_5	OR2AT4		GAGGTACATCTGCAGTAAGC	30665		26098
OR2AT4_6	OR2AT4		GCCAAGGTAGCATTGGTCTG	39134		23794
SLCO2B1_1	SLCO2B1		GGCCTGTACCTCGTTGAAGG	26629		31559
SLCO2B1_2	SLCO2B1		GCTGCTGGCCTCCTTCAACG	29576		136747
SLCO2B1_3	SLCO2B1		GGCCTCTCCAGCCAGACGTC	55532		9800
SLCO2B1_4	SLCO2B1		GGATATGCCACAGGACTTCA	0		12586
SLCO2B1_5	SLCO2B1		GTAGCTTGAGCAGTTGCCAT	53904		24573
SLCO2B1_6	SLCO2B1		GTTCGTGGCACAGACCCTGC	31161		148069
ARRB1_1	ARRB1			GGGAGAGGAGAGGCCATGGA	32637		45157
ARRB1_2	ARRB1			GAGTATCTCAAAGAGCGGAG	22521		20272
ARRB1_3	ARRB1			GCGTTCCTGCAGCCGCGTCA	33417		24052
ARRB1_4	ARRB1			GTCCCGCTTTCCCAGGTAGA	0		1
ARRB1_5	ARRB1			GTCCTCCCGGCCATAGCGGA	0		218
ARRB1_6	ARRB1			GGAACGCCTCATCAAGAAGC	10322		0
RPS3_1	RPS3			GGCCAAAGGCTGCGAGGTTG	29621		44271
RPS3_2	RPS3			GTGGTGTCTGGGAAACTCCG	2		5004
RPS3_3	RPS3			AGTTAACAGGGTCTCCGCTG	15056		8642
RPS3_4	RPS3			GCGCCACGTGTTGCTCAGAC	13404		15315
FLJ33790_1	FLJ33790		GGGCGGCGTCAACACGGACA	46481		115636
FLJ33790_2	FLJ33790		GGACCTAGCTGAAGTGATCG	51274		11862
FLJ33790_3	FLJ33790		GCCCTCGAGAAAGCGCACGC	10260		32936
GDPD5_1	GDPD5			GGCAAACACCACGGTGAAGA	71869		24224
GDPD5_2	GDPD5			GCTCCCGGGGCGGGTCACGC	10691		8744
GDPD5_3	GDPD5			GCATGGGCTACTGGAGCGAC	16752		12437
GDPD5_4	GDPD5			GGAACCAGAGGCGCTCCCAC	5257		589
GDPD5_5	GDPD5			GAGCGTGAGGCCAAAGGTGA	23926		3046
GDPD5_6	GDPD5			GTGGTCACAAGGATGGGTAC	28968		7086
SERPINH1_1	SERPINH1		GAACATCCTGGTGTCACCCG	42280		8516
SERPINH1_2	SERPINH1		GCCTGACATGCGTGACAAGT	15726		6104
SERPINH1_3	SERPINH1		GTGGTGGTGGCCTCGTCGCT	11709		6365
SERPINH1_4	SERPINH1		GGCGCTGCGCTCGGCAAGCG	5206		2335
SERPINH1_5	SERPINH1		GTCGCTAGGGCTCGTGTCGC	45211		1835
SERPINH1_6	SERPINH1		GAGGAGGTGCACGCCGGCCT	7797		11015
MAP6_1	MAP6			GGCGTGGCCGTGCATCACGA	28032		20799
MAP6_2	MAP6			GCCTACCGGCGAGCGCGAGC	21952		37733
MAP6_3	MAP6			GAGGGCTGGTATTCGCTGCG	15045		11988
MAP6_4	MAP6			GCATCCAACTCGCCCTGGGC	12952		18691
MAP6_5	MAP6			GATGCAGTTGCCCGGGCAAC	0		0
MAP6_6	MAP6			GCTGGTATTCGCTGCGCGGC	1		37044
MOGAT2_1	MOGAT2		GACCGAGACAAGCCACGGCA	34492		6006
MOGAT2_2	MOGAT2		GTCTGCAGCCTGCGCTCCCA	64135		22241
MOGAT2_3	MOGAT2		GCCCGCAATGTAGTTCCGAG	9298		79718
MOGAT2_4	MOGAT2		GAAGCCCACAGTGCAGATCT	21121		90404
MOGAT2_5	MOGAT2		GCTCCTCACTGTCCTGTATG	0		25063
MOGAT2_6	MOGAT2		GCCGGCACATCCAGGCCATC	1		0
DGAT2_1	DGAT2			GACCCCTCGCGCGACAGCGC	15954		25513
DGAT2_2	DGAT2			GACTGGAACACACCCAAGAA	26163		5329
DGAT2_3	DGAT2			GGCGGCTATGAGGGTCTTCA	57936		19746
DGAT2_4	DGAT2			GGACCTGCGCTGTCGCGCGA	3217		769
DGAT2_5	DGAT2			GCAGAATATGTACATGAGGA	10643		23811
DGAT2_6	DGAT2			GGTCACAGTGGGTCCGAAAC	0		996
UVRAG_1	UVRAG			GGGGCCCCCGACCGACGCGG	17533		9080
UVRAG_2	UVRAG			GCCTCCCGGTTCTGCCGCGC	14882		17614
UVRAG_3	UVRAG			GCTCATTTAGGGATTCCTTC	11313		71905
UVRAG_4	UVRAG			GAGACGGCAGCTCCACATGC	15117		62207
UVRAG_5	UVRAG			GAAGTGTAAAGTAGGTATCA	30004		11938
UVRAG_6	UVRAG			GATTGAATGGAAAGTCTGTT	39336		36974
WNT11_1	WNT11			GCTGCTTGCAGTGTTGCGTC	35591		32438
WNT11_2	WNT11			GGTGGTGCACGCCGCCCGCG	13968		49417
WNT11_3	WNT11			GCACTACCTTGGTGGCCGAC	30177		17716
WNT11_4	WNT11			GGCGCTGTCCAAGACACCAT	9094		24010
WNT11_5	WNT11			GGAGGGTCTGGTGTCTGCAC	16170		747
WNT11_6	WNT11			GGCCTTCATGACCTCGCGGG	1		880
PRKRIR_1	PRKRIR		GCGCTGCCCCCAACTGCACG	14725		37222
PRKRIR_2	PRKRIR		GTGCTCTTCCGCGTGCAGTT	26310		14177
PRKRIR_3	PRKRIR		GCTTACCTGGCAGGGTCCCG	5072		4999
C11orf30_1	C11orf30		GCACAGGGGGATCTCACCA	0		0
C11orf30_2	C11orf30		GACCTAATAGCTCTTCAGAA	56189		25381
C11orf30_3	C11orf30		GACCATTCTGAAGAGCTATT	17541		10407
C11orf30_4	C11orf30		GGAGAGCAGTAAACGATGAA	9959		5960
C11orf30_5	C11orf30		GTCCATTGAAGGTCGTCGAT	26582		12398
C11orf30_6	C11orf30		GGTTGACGTGGTACTTGTGT	5789		20347
LRRC32_1	LRRC32		GGACAAGAAGGTCTCGTGCC	19102		19095
LRRC32_2	LRRC32		GAGCATCTTACCATCTTACA	51408		99936
LRRC32_3	LRRC32		GACTGTTCTCCGCCAGTGAG	30089		114617
LRRC32_4	LRRC32		GTGTTGTGCAGCCAGGCCTA	24746		0
LRRC32_5	LRRC32		GCTGGTTCCCAGATAGATCA	20559		1305
LRRC32_6	LRRC32		GAAGTGCTGTGTAGAAGCCC	9298		30479
TSK_1		TSK			GAAGCTGTCGAAAAGGCCGA	11556		9929
TSK_2		TSK			GGACCTTTCGGGCACCAACC	13708		15504
TSK_3		TSK			GGGAACCCTCTAGCTGTCAT	25530		16829
TSK_4		TSK			GATGGGCACCGGCATGATGT	36225		26863
TSK_5		TSK			GGACCTGTCCTCCAACCGGC	33136		101214
TSK_6		TSK			GCGACTCCAGGTAGCGAAGG	15471		38989
PHCA_1	PHCA			GGACTGGGTTATACATCATT	21231		64648
PHCA_2	PHCA			ATGCAACAGCTGTATATCAT	1		19606
PHCA_3	PHCA			GTTCCTATTTAGGTCATGTA	3		11354
PHCA_4	PHCA			GATCTATTTATATTGTTACA	0		627
B3Gn-T6_1	B3Gn-T6		GAGGGCGGCTCGTCAGCCGC	18680		3215
B3Gn-T6_2	B3Gn-T6		GACTCTGGCCTGCCTCCTGG	55663		13771
B3Gn-T6_3	B3Gn-T6		TGACTGCCAAGACTCTGGCC	25490		30179
B3Gn-T6_4	B3Gn-T6		GGAGCGTCGGTGGGACCCTC	16600		48693
B3Gn-T6_5	B3Gn-T6		GCCGAGGCGTTCGCCACGCA	20441		7381
B3Gn-T6_6	B3Gn-T6		GCCGCCACTTCCCGCTGCTT	5		48153
CAPN5_1	CAPN5			GGGTCGCTTCCACCTGACGG	9147		21878
CAPN5_2	CAPN5			GAAGATGCCCGCGTAGGCGT	36375		50227
CAPN5_3	CAPN5			GGGCTTCACACACGAGAACA	56153		29075
CAPN5_4	CAPN5			GTAGAGTGAGTCGTCAGTGG	26445		33057
CAPN5_5	CAPN5			GATGCCATCCACAAAGAGGC	21222		61817
CAPN5_6	CAPN5			GGTGGGCAACTGCTGGTTTG	27417		80629
OMP_1		OMP			GTCGGCCTCATTCCAATCTA	39524		49546
OMP_2		OMP			GAGGCCGACGCCCTGGAGTT	14438		89191
OMP_3		OMP			GTGGAGAGCCTGAAGCAGCG	58149		15782
OMP_4		OMP			GTAGCCGCATCTGCCTGGTC	6630		7156
OMP_5		OMP			GAACTGTAGCCGCTGCTGCT	4846		1243
OMP_6		OMP			GTCCTCCTTGCGCCAGAAGA	57381		45296
MYO7A_1	MYO7A			GGATCATGTCCTCCACGCCG	5616		54526
MYO7A_2	MYO7A			GAGCTTCACCACCGCCCCGA	11326		6476
MYO7A_3	MYO7A			GACCTGCTGAAGAATCACCA	102589	7379
MYO7A_4	MYO7A			GAAGCTCTGCGACTCTGGGC	36618		12192
MYO7A_5	MYO7A			GTGGAGGACATGATCCGCCT	31911		39518
MYO7A_6	MYO7A			GCAACCTGCTTATCCGCTAC	35851		91704
GDPD4_1	GDPD4			GTCAAAGTTAAAGTATTCAC	30526		41570
GDPD4_2	GDPD4			GTGAATACTTTAACTTTGAC	19624		10818
GDPD4_3	GDPD4			GCAATGAAGAGTAGGCAGTC	14571		18556
GDPD4_4	GDPD4			GAAGAATCACTAGCAGAATC	22930		2894
GDPD4_5	GDPD4			GGTAGCTGGGCTGTCAATGC	11571		16955
GDPD4_6	GDPD4			GGCCTTCTACGTGGCATGTT	33083		14486
PAK1_1	PAK1			GATGCTGGAACCCTAAACCA	32561		1971
PAK1_2	PAK1			GAAAAACCCGCAGGCTGTTC	25851		27426
PAK1_3	PAK1			GCCGGCTCCAATCATAGTGC	0		3
PAK1_4	PAK1			GGTTCCAGCATCTTTGCTGC	0		0
PAK1_5	PAK1			GACATCAAATATCACTAAGT	25136		12205
PAK1_6	PAK1			GTGCTGGTATTTCTCATCGG	37664		9351
AQP11_1	AQP11			GTTACAGGAGGAAGTCTAAC	39754		64165
AQP11_2	AQP11			GGCTGGCCCGCGTAGTCGCC	15216		25718
AQP11_3	AQP11			GCTGTCGGTGGTGCTGCTCA	10425		0
AQP11_4	AQP11			GCTCAGCAGTTGCAGCTCGT	16021		27789
AQP11_5	AQP11			GTGCATGGCCTGACTCTGGT	38250		5656
CLNS1A_1	CLNS1A		GCCACACTGGAGAGATTAGA	43439		730
CLNS1A_2	CLNS1A		GCCGCTCTATCTGGCTAGCA	39610		34911
CLNS1A_3	CLNS1A		GCCAAATTTGAAGGTATGTA	29123		36482
CLNS1A_4	CLNS1A		GTACCCTTTACATCGCTGAG	31460		11120
CLNS1A_5	CLNS1A		GGATAATGCATGTAAACTAA	33113		85828
CLNS1A_6	CLNS1A		GCTCTCCTAGACAGTCACTT	93438		26241
HBXAP(RSF1)_1/HBXAP(RSF1)	GGACAGAAATATCATCACGG	45495		176724
HBXAP(RSF1)_2/HBXAP(RSF1)	GATGACATCAAAGAAGCCGA	23609		7767
HBXAP(RSF1)_3/HBXAP(RSF1)	GGTGCTGCAGGCGCCGCCGC	20968		17307
HBXAP(RSF1)_4/HBXAP(RSF1)	GCCGTTCCCTGAGCTGGAGC	18999		34083
HBXAP(RSF1)_5/HBXAP(RSF1)	GCCAAGAGTTTAACAGTACC	2		0
HBXAP(RSF1)_6/HBXAP(RSF1)	GCTCCTTCTTGGAGCGCTAC	5953		12157
PTD015_1	PTD015		GGTGTACAGACTCTTGTGAT	76079		22133
PTD015_2	PTD015		GATTGGCCGAGGGATGAGTG	6269		37880
PTD015_3	PTD015		GAAATTGCTTCCTTATCATG	4598		24885
KCTD14_1	KCTD14		GCTGGACTTCCGTGCTGTTA	21299		24549
KCTD14_2	KCTD14		GTTGTCTAGGACCGCCTTCC	35608		124396
KCTD14_3	KCTD14		GAAGCGGCCCTCCGCGTCCG	5261		28218
KCTD14_4	KCTD14		GGGTACCCTGAGGAAGTTTC	76188		20820
KCTD14_5	KCTD14		GATGGGTCTGAAATAGGTGC	8516		1987
KCTD14_6	KCTD14		GAGCCTCACGGTACACTTCA	27447		21886
THRSP_1	THRSP			GAACTGCCTGCTGACCGTCA	52107		1519
THRSP_2	THRSP			GCCTGCTGACCGTCATGGAC	37775		5012
THRSP_3	THRSP			GGCTGCTGCCGCGGGAGGAG	22685		2254
THRSP_4	THRSP			GTGAAGTAGGTGTAGAGATC	88251		61300
THRSP_5	THRSP			GAATGGAACCGCAGAGACAG	28221		524
THRSP_6	THRSP			GGACCATGGGCTGCTGCCGC	15288		24987
NDUFC2_1	NDUFC2		GCGGCGCTCACAATTGCCAC	12757		57079
NDUFC2_2	NDUFC2		GAGCCGCGGGTCGGTCAGCT	18869		23214
NDUFC2_3	NDUFC2		GCGTCACCATGATCGCACGG	4753		29977
NDUFC2_4	NDUFC2		GCGGGTCGGTCAGCTTGGGC	10279		31098
NDUFC2_5	NDUFC2		GGCTTCTTGGGCTACTGCTC	10715		18825
NDUFC2_6	NDUFC2		GATGTAGAGGAGCCGCGGGT	0		616
ALG8_1	ALG8			GAGGGAATAATGTCCTGTTG	33455		16342
ALG8_2	ALG8			GTTGCCAAGGGAGTCACTGA	47265		8080
ALG8_3	ALG8			GGCGGCGCTCACAATTGCCA	89109		12389
ALG8_4	ALG8			GGCAACTTCAGAGTGGACGT	0		5855
ALG8_5	ALG8			GTGCGAGTCCCTTACTATGT	41524		43083
ALG8_6	ALG8			GTATTACTTCTGTGGAACTT	99009		145879
LOC283219_1	LOC283219		GTACCTCTTCCACGAAGACC	9561		42950
LOC283219_2	LOC283219		GACCCCATCACGCTGAACGT	57344		40753
USP35_1	USP35			GGAGCCACCATCTAGCGCCC	56618		38319
USP35_2	USP35			GGCACCAACCTTCTCAATTT	4997		5162
USP35_3	USP35			GGACAAGATCTTGGAGGCGG	19157		41766
USP35_4	USP35			GGCGCGCGCCTCTACGTGGG	23609		8086
USP35_5	USP35			GAAGAACTCGGCGAAGACGT	5188		2653
USP35_6	USP35			AGGCACTGCTCACGCTCCAG	5178		3959
GAB2_1	GAB2			GGTTCAAACGTGAACAGAGT	24616		29823
GAB2_2	GAB2			GATGATCCGCAGAGGCTTCT	42674		6218
GAB2_3	GAB2			GAACATCCATATTGTCTACC	23234		41919
GAB2_4	GAB2			GCTCCTTCTTGTTAAAGGTC	0		7553
GAB2_5	GAB2			GGCTGAGGAAACATTTCTCA	70953		110244
GAB2_6	GAB2			GGGCTGAGGACTTGCGCTCT	1		35599
dummy1_1	dummy1		GGTCTCTGTACGGGCCGCCC	45250		40053
dummy1_2	dummy1		GTGTCGGATTCCGCCGCTTA	19785		23063
dummy1_3	dummy1		GAATCGACCGACACTAATGT	43648		61988
dummy2_1	dummy2		GACTTCTAGAATATAAAAGA	13592		2289
dummy2_2	dummy2		GATGGCGCTTCAGTCGTCGG	23176		11429
dummy2_3	dummy2		GGTTAGAGACTAGGCGCGCG	26735		124049
dummy3_1	dummy3		GAGAAGGATGGAAATTAGAA	15999		5597
dummy3_2	dummy3		GCCGTGTTGCTGGATACGCC	38411		161246
dummy3_3	dummy3		GCGAACCCCGTAGCCAGGCT	72518		52186
dummy4_1	dummy4		GTAGGCGCGCCGCTCTCTAC	46625		29147
dummy4_2	dummy4		GTTCGCTTCGTAACGAGGAA	39804		54527
dummy4_3	dummy4		GTACCATACCGCGTACCCTT	26957		6126
dummy5_1	dummy5		GGGCCCGCATAGGATATCGC	19592		31340
dummy5_2	dummy5		GTCGTCCGGGATTACAAAAT	41215		130409
dummy5_3	dummy5		GTCATCAGCGATTTGACGAG	37753		19152
PLK1_HEG_1	PLK1_HEG		GCTCCGGGAGCTGCAACTCC	63874		20832
PLK1_HEG_2	PLK1_HEG		GCCGGCTGCGTGGGTCCACT	0		32784
PLK1_HEG_3	PLK1_HEG		GTCCGAGATCTCGAAGCACT	24115		6832
PLK1_HEG_4	PLK1_HEG		GCTGCTCAAGCCGCACCAGA	22014		4105
PLK1_HEG_5	PLK1_HEG		GAATCCTACGACGTGCTGGT	106813	5004
PLK1_HEG_6	PLK1_HEG		GTTGGAGCTCTGCCGCCGGA	5654		1
ACLY_HEG_1	ACLY_HEG		GAACCGATTCTGGATGGCTG	0		1
ACLY_HEG_2	ACLY_HEG		GGGTCACTCCTGACACAGAC	26752		155817
ACLY_HEG_3	ACLY_HEG		GACCAGCTGATCAAACGTCG	0		4770
ACLY_HEG_4	ACLY_HEG		GTCCTGGCTGAAGCCACGGC	66065		4673
ACLY_HEG_5	ACLY_HEG		GCCTCACACATACCTGACTG	100147	19639
ACLY_HEG_6	ACLY_HEG		GCATCTATGCCACCCGAGAA	43332		16960
AURKA_HEG_1	AURKA_HEG		GACACAAGACCCGCTGAGCC	21096		8703
AURKA_HEG_2	AURKA_HEG		GTGCTTGCAAAGGAATGCGC	37149		11763
AURKA_HEG_3	AURKA_HEG		GCTTGTCTCCAGTCACAAGC	17979		4115
AURKA_HEG_4	AURKA_HEG		GGAGACAGGATGAGGTACAC	9325		5514
AURKA_HEG_5	AURKA_HEG		GTTTACCAGGTGCCGATGGC	58259		35099
AURKA_HEG_6	AURKA_HEG		GCTTTGGAAGACTTTGAAAT	36758		28208
CCNH_HEG_1	CCNH_HEG		GTATCTAGTCCTCAGTTTGT	75155		63450
CCNH_HEG_2	CCNH_HEG		GAGTCCTCTTGGACAGGAGA	11656		55789
CCNH_HEG_3	CCNH_HEG		GAAGGCACTTGAACAGATAC	1		86334
CCNH_HEG_4	CCNH_HEG		GGGCTTCCTCATCGACTTAA	22441		34563
CCNH_HEG_5	CCNH_HEG		GTCCAAGAGGACTCTCCCGG	44833		3414
CCNH_HEG_6	CCNH_HEG		GTAGACCCGCTATCCCATAT	16413		5385
CDK2_HEG_1	CDK2_HEG		GGTCAATCTCAGAATCTCCA	42631		63998
CDK2_HEG_2	CDK2_HEG		GGTGGAGGACCCGATGAGAA	27429		1510
CDK2_HEG_3	CDK2_HEG		GAGAAGCATTACCTTGATGA	24374		11989
CDK2_HEG_4	CDK2_HEG		GAGGTTTAAGGTCTCGGTGG	15022		9211
CDK2_HEG_5	CDK2_HEG		GTGTTAATAAGCAGATTCTG	45071		4159
CDK2_HEG_6	CDK2_HEG		GCCATCAAGCTAGCAGACTT	0		0
```

**Supplemental Table 2. IC_50_ Values of P4HA3 and SERPINH1 against IntClust-2 and non-IntClust-2 cell lines.** The results are expressed as mean values out of three independent experiments at 48 h. ND: not determined.

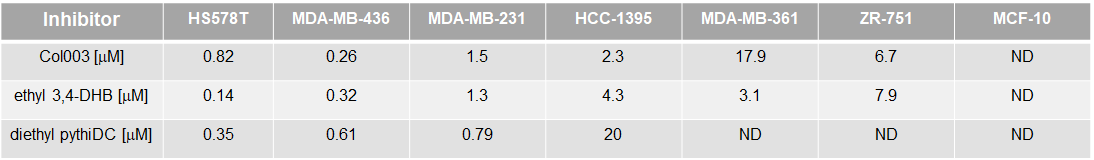

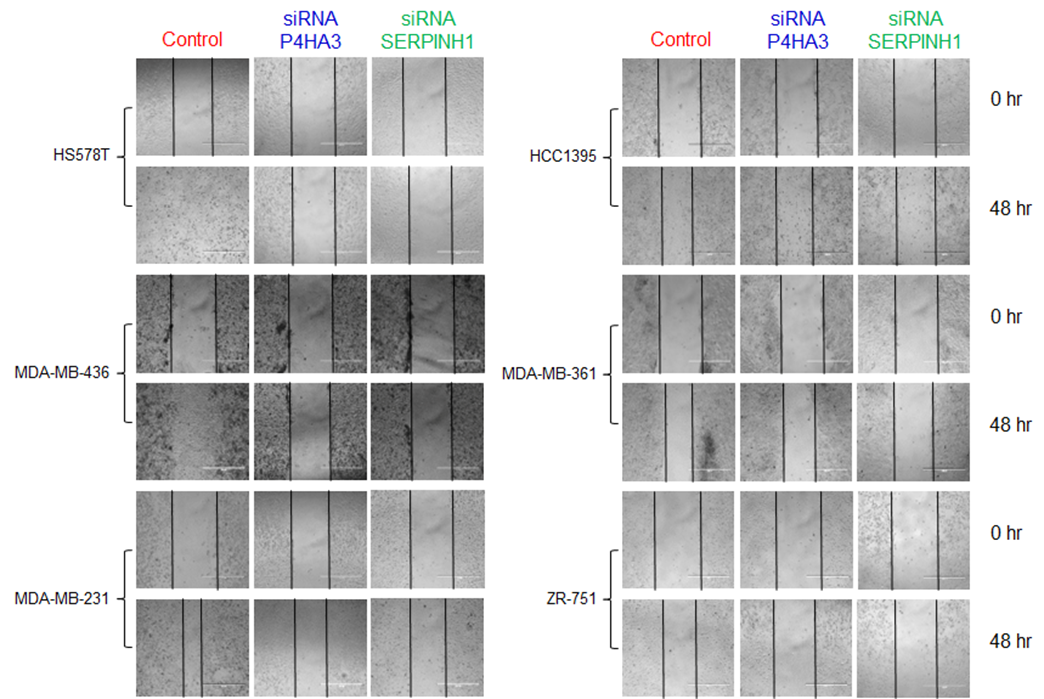

**Supplemental Figure 1. siRNA-mediated wound healing assay.** IntClust-2 and non-IntClust-2 cell lines were treated with the respective siRNA (25 nM). Images of the scratched areas were taken at 0 and 48 h using an EVOS fluorescence microscope.

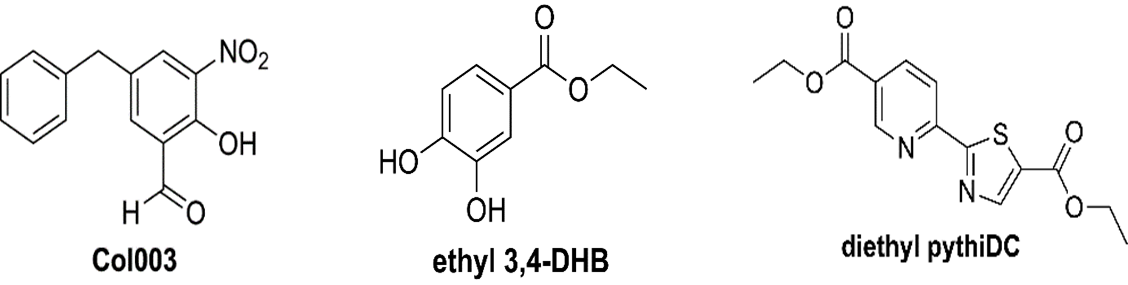

**Supplemental Figure 2. Chemical structures of the three key compounds.**

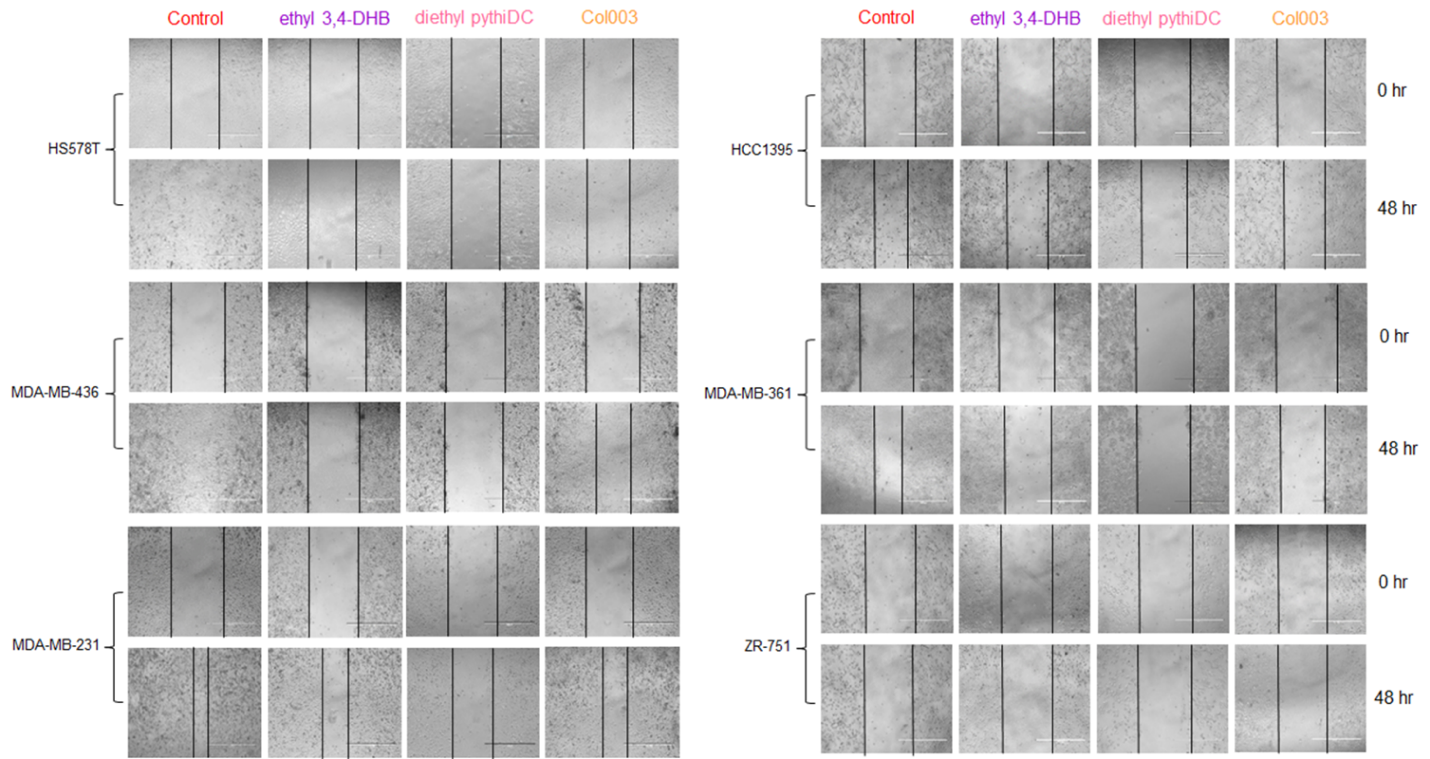
**Supplemental Figure 3. Drug-mediated wound healing assay.** IntClust-2 and non-IntClust-2 cell lines were treated with the respective small molecule inhibitors (2 mM). Images of the scratched areas were taken at 0 and 48 h using an EVOS fluorescence microscope.
